# Transcriptomics and trans-organellar complementation reveal a limited signaling capacity of 12-*cis*-oxo-phytodienoic acid in wounded Arabidopsis

**DOI:** 10.1101/2024.03.22.586262

**Authors:** Khansa Mekkaoui, Ranjit Baral, Fiona Smith, Moritz Klein, Ivo Feussner, Bettina Hause

## Abstract

12-*cis*-Oxo-phytodienoic acid (OPDA), the main precursor of the key plant growth and defense hormone jasmonoyl-isoleucine (JA-Ile), is believed to have distinct signaling roles in plant’s responses to stress. In Arabidopsis, insights into OPDA functions have been gained from studying a mutant, which is affected in the conversion of OPDA by missing OPDA REDUCTASE3 (OPR3). *opr3* mutants, however, accumulate JA-Ile through a cytosolic bypass mediated by OPR2. Therefore, wound-induced transcriptome of *opr2opr3* in comparison to wild-type and *allene oxide synthase* mutant was analyzed to unravel OPDA signaling. Results showed that OPDA lacked a distinct transcriptional signature, whereas known OPDA-response genes were wound-induced independently of OPDA. The application of OPDA to *opr2opr3* resulted in a distinct transcriptional response compared to the endogenous rise of OPDA in the same mutant, with the activation of the sulfur assimilation pathway genes occurring only with the external application of the compound. These findings suggested a compartmentalization of endogenously produced OPDA, investigated further through trans-organellar complementation. OPR3 complemented *opr2opr3* mutants in fertility and wound-induced JA-Ile production regardless of its localization. Since *in vitro* enzymatic studies revealed OPR3’s activity on both OPDA and 4,5-ddh-JA, conclusions on translocation of OPDA were not unequivocal. Dissecting the conversion of either OPDA or 4,5-ddh-JA by OPR2 and OPR1 organelle variants pointed, however, to a strong OPDA compartmentalization supporting its lacking signaling function.

## Introduction

Jasmonic acid (JA) and its bioactive form jasmonoyl-isoleucine (JA-Ile) function primarily as signaling molecules coordinating plant growth and reproduction and responses to biotic and abiotic stresses ^1–4^. They play a central role in the defense of plants against mechanical damage and herbivory, as JA-Ile rapidly accumulates at the site of injury and acts as a signaling molecule by triggering a cascade of defense responses ^5,6^. JA-Ile signaling involves a series of molecular events mediated by its perception by the F-box protein CORONATINE INSENSITIVE 1 (COI1) leading to the degradation of the JASMONATE ZIM-DOMAIN (JAZ) repressors, hence the release of the expression of JA-responsive genes ^7–13^.

One major intermediate in the formation of the bioactive JA-Ile, 12-*cis*-oxo-phytodienoic acid (OPDA), has been known to be a signaling molecule with independent functions from JA-Ile ^14–17^. OPDA formation takes place in the chloroplast via the octadecanoid pathway, by oxygenation of α-linolenic acid by 13-lipoxygenases (LOXs) and the coupled actions of ALLENE OXIDE SYNTHASE (AOS) and ALLENE OXIDE CYCLASE (AOC) ^18–20^ (Fig. 1a). Metabolization of OPDA into JA occurs after its transfer to peroxisomes, where the reduction of its cyclopentenone ring by OPDA REDUCTASE3 (OPR3) and three rounds of β-oxidation take place ^1,21,22^. A parallel hexadecanoid pathway gives rise to *dinor*-12-oxo-phytodienoic acid (*dn*-OPDA), which in turn is converted to JA following the same pathway as OPDA, but with two rounds of β-oxidation only ^23^.

**Fig. 1:**
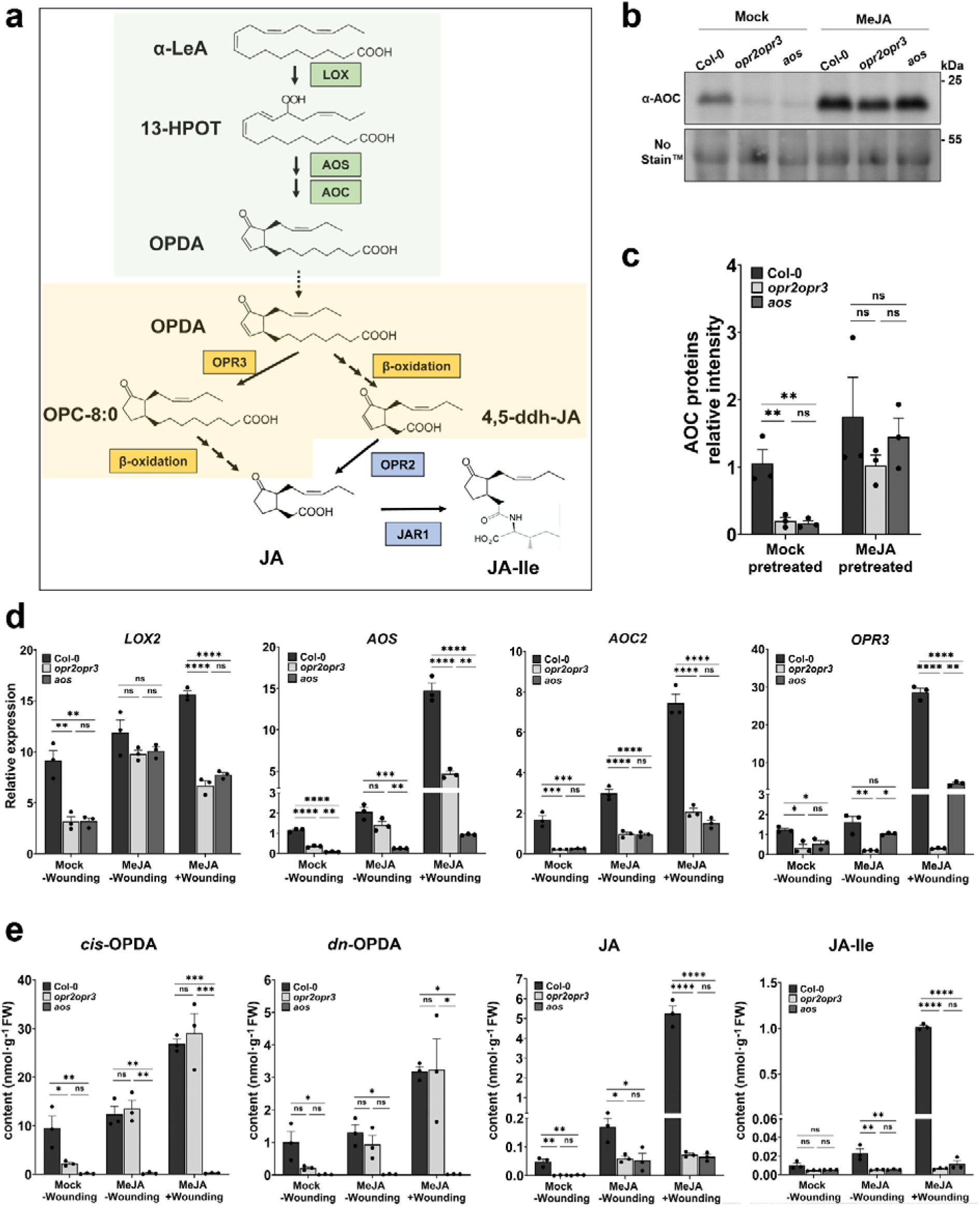
MeJA pretreatment restitutes AOC protein levels in the mutants and OPDA content in the *opr2opr3* mutant. **(a)** Synthesis of jasmonic acid (JA)/JA-Ile from α-linolenic acid (α-LeA). The biosynthesis starts with the release of α-LeA from chloroplast membrane galactolipids and its conversion to (13S)-hydroperoxyoctadecatrienoic acid (13-HPOT) by 13-LIPOXYGENASE (LOX). *cis*-(+)-12-oxophytodienoic acid (OPDA) is formed in the chloroplast (green background) from 13-HPOT through sequential actions of ALLENE OXIDE SYNTHASE (AOS) and ALLENE OXIDE CYCLASE (AOC). In the peroxisomes (yellow background) reduction of cyclopentenone ring of OPDA is catalyzed by a peroxisomal OPDA REDUCTASE (OPR3) resulting in the formation of 3-oxo-2-(20(Z)-pentenyl)-cyclopentane-1-octanoic (OPC-8:0). Shortening of the octanoic acid side chains of OPC-8:0 by three rounds of β-oxidation yields JA. The cytosol-released JA is conjugated to Isoleucine (Ile) by the enzyme JASMONATE RESISTANT 1 (JAR1) resulting in the major bulk formation of the biologically active JA-Ile. OPDA can directly undergo three rounds of β-oxidation yielding 4,5-didehydro-JA (4,5-ddh-JA) in a parallel OPR3-bypass pathway. The formed 4,5-ddh-JA is released to the cytosol and then reduced to JA by OPR2 resulting in JA-Ile formation. Enzymes shaded in green, yellow and blue are located in the plastids, peroxisomes and cytosol, respectively. **(b)** AOC protein content in seedlings pretreated with 1 µM MeJA or water (Mock) visualized by immunoblot, equal loading is depicted by staining with No-Stain^TM^ Protein Labeling Reagent. **(c)** AOC band intensity quantified relative to total protein loading. **(d)** RT-qPCR of JA-responsive genes *LOX2*, *AOS*, *AOC2* and *OPR3* showing transcript accumulation at control condition (-wounding) and at 1 h after wounding. Seedlings were pretreated with MeJA or water (Mock) during development. **(e)** Levels of *cis*-OPDA, *dn*-OPDA, JA and JA-Ile in seedlings of wild-type (Col-0), *opr2opr3*, and *aos* at control condition (-wounding) and 1 h after wounding. For detailed values see Supplementary Tab. 2. Seedlings were pretreated with MeJA or water (Mock) during development (see Supplementary Fig. 2a for experimental set-up). Bars represent means of three biological replicates with 120 seedlings each (single dots; ±SEM). Statistically significant differences among genotypes within each condition were calculated using One-Way ANOVA followed by Tukey-HSD and are indicated by asterisks with *p<0.05, **p<0.01, ***p<0.001, and ****p<0.0001.

The putative distinct function of OPDA in Angiosperms partially derives from its function in non-vascular plants by inducing defense- and growth inhibition-related transcriptional reprogramming in *Marchantia polymorpha* ^13,24–26^. Such JA-independent functions of OPDA were also characterized in vascular plants for different vegetative processes including initiation of tendril coiling in *Bryonia dioica* ^27^, regulation of embryo development in *Solanum lycopersicum* ^28^, and inhibition of seed germination and induction of stomatal closure in *Arabidopsis thaliana* ^29–31^. Despite its perception being not identified in Arabidopsis, OPDA was shown to act as signaling molecule by transcriptionally inducing a set of genes, which is not induced by JA ^32^. These OPDA-specific response genes were characterized as wound- and OPDA-responsive in a COI1-independent manner and encompassed mainly stress-responsive genes, e.g. *DRE-BINDING PROTEIN 2A* (*DREB2A)*, *FAD-LINKED OXIDOREDUCTASE* (*FAD-OXR), SALT TOLERANCE ZINC FINGER* (*ZAT10)*, *ETHYLENE RESPONIVE ELEMENT BINDING FACTOR5* (*ERF5*), *GLUTATHION TRANSFERASE6* (*GST6*), and the glutaredoxin member *GRX480*. OPDA contains an α, β-unsaturated carbonyl group which is thought to react with nucleophilic molecules by Michael addition ^33^. The electrophilic nature of oxylipins could convey biological activities, as deduced from the accumulation of oxylipins adducts, including OPDA-GSH conjugates, during the hypersensitive response in pathogen-elicited tobacco leaves ^34^. OPDA binding to the CYCLOPHILIN20-3 (CYP20-3) was suggested to trigger the formation of the cysteine synthase complex activating sulfur assimilation to regulate cell redox homeostasis through mediation of *TGACG-BINDING* (TGA) transcription factors ^35,36^. Here, CYP20-3-dependent OPDA signaling induces high levels of GSH leading to a retrograde signaling, which coordinates expression of OPDA-induced genes ^37^. These transcriptional changes in Arabidopsis induced by OPDA through CYP20-3 module and/or additional uncharacterized mechanism consolidated the hypothesis of its function not limited to being only a JA precursor ^16^.

The previous work characterizing OPDA signaling and JA-independent functions in Arabidopsis relied mainly on the use of the *opr3* mutant as the genetic background where OPDA and JA biosynthesis were uncoupled due to missing OPR3 ^38,39^. The *opr3* mutant is male sterile like the *aos* mutant being deficient in OPDA and JA ^38,40,41^. Both mutants show defects in anther dehiscence, stamen elongation and pollen development showing that these defects were due to impairments in JA production. Persistent resistance of the *opr3* mutant against necrotrophic pathogens initially hinted towards a defense function of OPDA, but further studies uncovered a conditional removal of the T-DNA containing intron upon fungal infection of *opr3-1* on the one hand ^42^, and an OPR3-independent pathway leading to JA/JA-Ile formation through the cytosolic OPR2 in the *opr3-3* mutant on the other hand ^43^. Low JA-Ile formation in both *opr3* mutants rather than OPDA was shown to be responsible of its resistance, hence challenging the suggested JA-independent OPDA signaling function ^42,43^. Mainly the recently identified formation of JA-Ile via bypassing the OPR3 (Fig. 1a) questioned previously identified functions of OPDA. Consequently, OPDA-specific signaling exclusively analyzed with OPDA-producing but JA/JA-Ile deficient mutants requires further studies.

This study re-addressed the question whether OPDA can function as an independent signaling molecule in Arabidopsis besides being a JA precursor. Using the *opr2-1 opr3-3* mutant (further named *opr2opr3*) where formation of JA/JA-Ile does not occur ^43^, we investigated the effects of basal and wound-induced OPDA on gene expression in seedlings and rosette leaves of *A. thaliana*. The results suggested that the endogenously formed OPDA is unlikely to be responsible for mediating gene expression, while exogenous supply of OPDA resulted in a distinct signaling. The activation of the sulfur assimilation pathway exclusively by exogenously supplied OPDA hints towards its predominant detoxification within the cell but indicates a compartmentalization of the endogenously formed OPDA. Therefore, we addressed the question of the occurrence of OPDA within the cell and whether it could be perceived. A trans-organellar complementation approach showed that OPDA is likely to be confined to the cell compartments where it is synthesized (plastids), transported through (cytosol) and metabolized (peroxisomes). Instead, 4,5-ddh-JA appears to be a mobile compound within the plant cell supporting the assumption that OPDA has no signaling function in wounded Arabidopsis plants.

## Results

### A JA positive feedback loop during development is responsible for the tissue capability to produce OPDA

The *opr2opr3* double mutant was characterized as JA-deficient, with JA formation through the peroxisomal OPR3 and alternatively through the cytosolic OPR2 being both interrupted ^43^. The JA deficiency in the *opr2opr3* and *aos* mutants was confirmed by measuring JA content under both control and wounding conditions (Supplementary Fig. 1a). In the mutants, JA levels remained at the detection limit, with approximately 0.002 nmol/g of fresh weight JA detected in the wounded *opr2opr3*. This contrasts with the reported 0.2 nmol/g of fresh weight JA accumulation in the wounded *opr3* mutant plants^39^, thereby confirming the JA deficiency of *opr2opr3* ^43^. JA deficiency is associated with a decrease in the accumulation of JA-biosynthesis enzymes, among them specifically the AOC, due to the disruption of the JA positive feedback loop during development ^44^. Consequently, lower background and wound-induced levels of OPDA were observed in the *opr2opr3* mutant compared to the wild-type (Supplementary Figure 1b), which indeed correlated with diminished AOC protein content in *opr2opr3* and *aos* seedlings in comparison to wild-type (Col-0) seedlings (Supplementary Fig. 1c, d).

Due to the partial OPDA deficiency of *opr2opr3* seedlings, it was crucial to normalize the levels with those of the wild type prior to investigating OPDA signaling within this mutant. To achieve this, we mimicked the missing JA positive feedback loop in the *opr2opr3* and *aos* mutants by supplying seedlings with 1 µM JA methyl ester (MeJA) during development to increase the AOC protein levels (Supplementary Fig. 2a). In contrast to water (mock-)pretreated seedlings, seedlings pretreated with MeJA during development exhibited increased AOC protein levels in wild-type and both mutants (Fig. 1b). Notably, this increase was more pronounced in the mutants resulting in AOC protein levels in *opr2op3* and *aos* seedlings matching those of wild type seedlings (Fig. 1c). Transcript accumulation of the JA biosynthesis enzymes *LOX2*, *AOS*, *AOC2* and *OPR3* determined by RT-qPCR corroborated the function of the JA positive feedback loop: MeJA pretreatment during development resulted in either a compensation or an enhancement of their basal transcript levels in the mutants compared to wild type (Fig. 1d). They all are, however, highly induced upon wounding in wild-type seedlings, but their induction is significantly lower in the mutants (Fig. 1d). This dampened induction demonstrates that the MeJA pretreatment did not activate a wound-induced JA-Ile signaling in the JA-deficient mutants. Most importantly, however, the restoration of AOC protein content in the *opr2op3* enhanced its basal and wound-induced OPDA and *dn*-OPDA levels to the levels of wild-type, whereas the *aos* mutant remained OPDA deficient (Fig. 1e). Despite the accumulation of low residual levels of JA in *opr2opr3* and *aos* seedlings upon pretreatment, wound-induced accumulation of JA and JA-Ile was only observed in wild-type seedlings and correlated with the transcriptional response of the JA-responsive genes (Fig. 1d). This validates the JA deficiency of both mutant lines within this experimental design and the exclusive OPDA enhancing role of the developmental JA-feedback loop for the *opr2opr3* mutant (Fig. 1e).

### A comparative transcriptomics approach does not show an OPDA-specific response to wounding

OPDA signaling in Arabidopsis was previously inferred from its capacity to mediate transcriptional changes by inducing a specific set of OPDA-responsive genes in the *opr3-1* single mutant ^32,45^. Another putative OPDA-signaling pathway was based on the detected binding of OPDA to CYP20-3, which in turn transcriptionally induces TGA transcription factors ^35^. To further clarify OPDA-related signaling function, a comparative transcriptomics approach was set between wild-type and *opr2opr3* seedlings. To uncouple putative OPDA signaling from common wound-induced signaling processes, *aos* seedlings were included as a negative control, since both JA and OPDA production are abolished in this mutant (Supplementary Fig. 2b). mRNA sequencing of MeJA-pretreated seedlings from Col-0, *opr2opr3* and *aos* was performed using unwounded seedlings (control) and seedlings harvested 1 h after wounding. The overall wound-induced response showed a significant transcriptional change in all three genotypes (Supplementary Fig. 3). Here, a strong similarity between the transcriptomes of *opr2opr3* and *aos* seedlings was evident in the expression heatmap, with a clear overlap observed in the principal component analysis (PCA), irrespective of the condition (Supplementary Fig. 3). The wounding transcriptional response showed most difference in the number of differentially expressed genes (DEGs) between Col-0 and the JA-deficient mutants, with the DEGs being 10.84% and 7.48% greater in Col-0 than *opr2opr3* and *aos*, respectively (Fig. 2a). A common wound response was observed in all the genotypes which was enriched in genes involved in stress, oxygen, and abiotic stimulus responses and was therefore rather wound-inducible but completely independent from the JA pathway (Fig. 2a, b, Supplementary Dataset 1). Interestingly, this group contained known stress-responsive genes with several genes being previously reported as OPDA-responsive, such as *DREB2A* and *FAD-OXR,* which were also induced in the *aos* mutant by wounding despite its OPDA deficiency^32^ (Fig. 2c, d, Supplementary Tab. 4, Supplementary Dataset 1). Additionally, other major OPDA-marker genes like *ZAT10, ERF5, TCH4* and *GST6* also showed a wound-responsive induction in *aos* seedlings indicating that they are more likely to be associated with an OPDA-independent wound-induced stress response rather than with an OPDA-mediated response (Fig. 2d). To exclude a JA/JA-Ile dependence of their induction, transcript levels of these genes were monitored in rosette leaves of two *coi1*-mutant lines, *coi1-16* and *coi1-30* (Supplementary Fig. 4a, b). The data show that all tested genes are inducible by wounding to nearly the same level as in wild type, irrespective of the lower levels of OPDA in the *coi1*-mutants reaching in *coi1-16* only 25 % of that of wounded wild type leaves, whereas there was no increase in wounded leaves of *coi1-30* (Supplementary Fig. 4c). Hence, their transcriptional induction by wounding did not correlate with OPDA levels despite its COI1-independence indicating that they are rather governed by other signaling processes in the wound response. It is interesting to note that also JA and JA-Ile levels appeared to be significantly lower in wounded *coi1* plants in comparison to wild type pointing again to the missing positive feedback in JA signaling and biosynthesis during plant development ^44^.

**Fig. 2:**
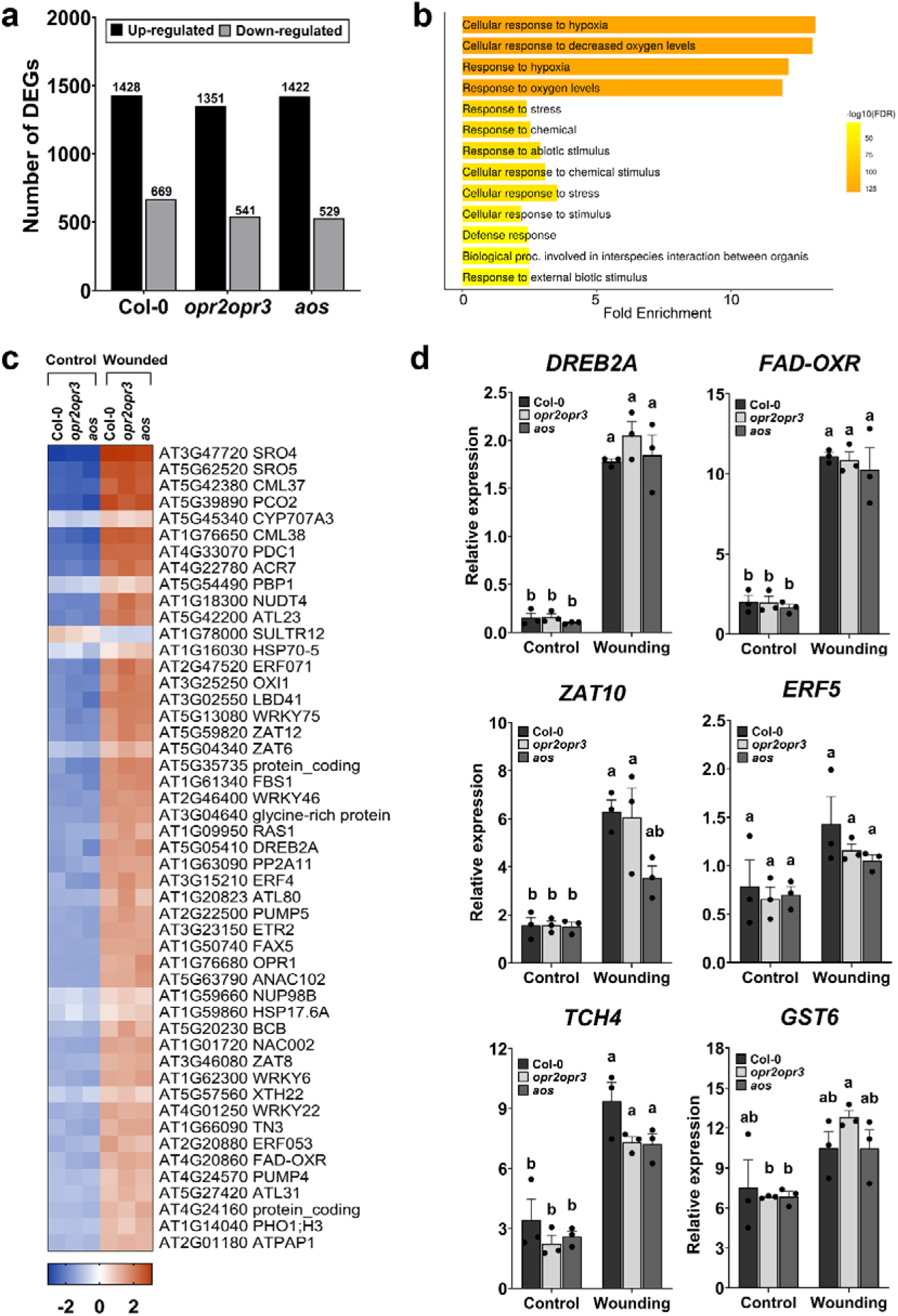
The JA/OPDA-independent wound response of seedlings. Ten-day old seedlings from Col-0, *opr2opr3* and *aos* were pretreated with MeJA during development and were either unwounded (control) or harvested 1 h post wounding with forceps (wounding). **(a)** Number of differentially regulated genes (DEGs) after wounding compared to non-wounded controls in seedlings of Col-0, *opr2opr3* and *aos*, all pretreated with 1 µM MeJA during development (see Fig. S2 for experimental set-up). DEGs were identified using FDR and FC cutoffs of 0.05 and 2, respectively. **(b)** Bar plot showing Gene Ontology enrichment analysis summarizing the significant biological processes (FDR=0.05) enriched commonly in all genotypes in response to wounding, with bars indicating gene fold enrichment and color scale indicating FDR values. **(c)** Heatmap illustrating 50 out of 211 DEGs enriched in the GO term ‘response to stress’ from **(b)**. Average FPKM values of three independent biological replicates were transformed into row normalized Z-score. **(d)** RT-qPCR validation of DEGs previously described as OPDA-responsive genes. Transcript accumulation of *DREB2A*, *FAD-OXR*, *ZAT10, ERF5, TCH4,* and *GST6* in Col-0, *opr2opr3* and *aos* at control and wounding conditions. Transcript levels were normalized to those of *PP2A3*. Bars represent means of three biological replicates (single dots; ±SEM). Statistically significant differences among genotypes within each condition were calculated using Two-Way ANOVA followed by Tukey-HSD and are indicated by different letters.

In wild-type seedlings, the signaling mediated by JA/JA-Ile was evident as it distinctly shaped the transcriptome, sustaining statistically differential basal levels of 71 JA-regulated genes when directly comparing the wild-type transcriptome to those of the mutants at control condition (Supplementary Fig. 5a, b). Wounding of wild-type seedlings resulted in a more drastic transcriptional change highlighted by 432 DEGs specifically induced in Col-0 and enriched in genes related to JA, wounding, and immune responses (Supplementary Fig. 5a, c, Supplementary Dataset 2). These transcriptional responses, notably of genes encoding JAZ transcription factors and other JA-responsive genes, such as *CHLOROPHYLLASE 1* (*CLH1)* and *N-ACETYLTRANSFERASE ACTIVITY 1* (*NATA1)*, were dampened in the *opr2opr3* and *aos* mutants confirming the JA dependence of their induction upon wounding (Supplementary Fig. 5d, Supplementary Dataset 2). These data suggest that next to a common wound response, JA/JA-Ile are main mediators of wound-induced transcriptional changes in wild-type seedlings.

### Wound-induced and exogenously applied OPDA resulted in different alterations of the transcriptome in *opr2opr3* seedlings

To identify OPDA-regulated genes, transcriptional differences between *opr2opr3* and *aos* were dissected and indicated a restricted number of DEGs at control and wounding conditions (Fig. 3a). Except for the mutated genes, which showed diminished expression in the respective mutant backgrounds (*AOS*, *OPR2*, and *OPR3*), only nine specific genes exhibited differential expression in *opr2opr3* seedlings compared to *aos* seedlings, regardless of the experimental condition (Fig. 3b, c). These genes encode five RNA species and four proteins, among them one cyclic nucleotide-gated ion channel (*CNGC11*), one cytochrome P450 (*CYP81D11*), and two hypothetical proteins. This is a very narrow or almost no difference between the transcriptomic profiles of the mutants, despite the presence of significantly high OPDA levels in the *opr2opr3* mutant compared to its absence in *aos* (Fig. 1e). These results make an OPDA-mediated gene expression in wounded seedlings of the *opr2opr3* mutant unlikely.

**Fig. 3:**
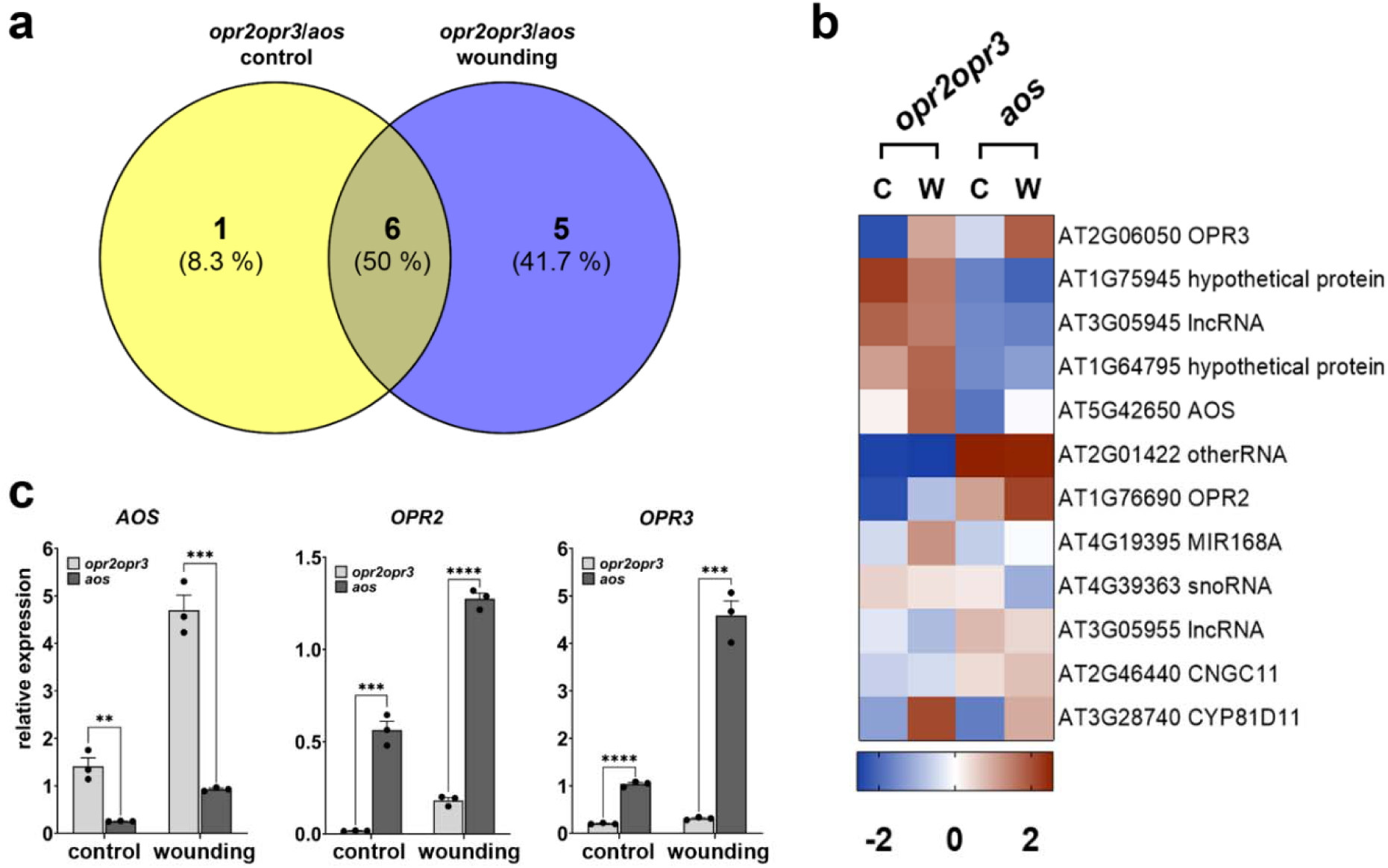
OPDA-specific transcriptional change deduced from transcriptomic comparison between seedlings of *opr2opr3* and *aos* at control and wounding conditions. Ten-day old seedlings from *opr2opr3* and *aos* were pretreated with MeJA during development and were either unwounded (control, C) or harvested 1 h post wounding with forceps (wounding, W). **(a)** Venn diagram representing genes showing differential expression when *opr2opr3* is compared to *aos* at control and wounding conditions (FDR cutoff = 0.05 and FC cutoff = 2). **(b)** Heatmap illustrating the expression levels of the eleven wounding-related DEGs from (A). The average FPKM values of three independent biological replicates were transformed into row-normalized Z-score. **(c)** Transcript accumulation of *AOS*, *OPR2,* and *OPR3* selected from **(b)** and determined by RT-qPCR. Transcript levels were normalized to those of *PP2A3*. Bars represent means of three biological replicates (single dots; ±SEM). Asterisks denote statistically significant differences among genotypes as determined by Student’s t-test within each condition (** p<0.01, *** p<0.001, **** p<0.0001).

To investigate the impact of developmental stage on our findings, the wounding experiment was done using mature four-week-old rosettes (Supplementary Fig. 2c). Pretreatment with 1 µM MeJA enhanced OPDA levels in wounded leaves of *opr2opr3* compared to the mock pretreated ones (Supplementary Fig. 6a). Regardless of whether the MeJA pretreatment was administrated, wounding led to a high number of DEGs in the wild-type and a lower, but similar number of DEGs in the *opr2opr3* and *aos* mutants (Supplementary Fig. 6b, c). Most importantly, consistent with the findings for seedlings, the transcriptomes of the *aos* and *opr2opr3* mutants were highly similar despite significant OPDA accumulation in the latter and differed only with the same DEGs found in the transcriptomes of seedlings (Supplementary Fig. 6d-f).

Previous work demonstrated a signaling role of OPDA in Arabidopsis by its exogenous supply to wild-type plants or to the *opr3* mutant ^32,35,45^. To determine whether the origin of OPDA caused the different outcomes, RNAseq data resulting from wounding of *opr2opr3* seedlings (= endogenous rise of OPDA) were compared to transcriptomic data generated from *opr2opr3* seedlings, which were treated with 25 µM OPDA for 30 min (= exogenous supply of OPDA). OPDA treatment resulted in transcriptional changes that were highly different from those induced by wounding (Supplementary Fig. 7a, Supplementary Dataset 3). The clustering of DEGs induced by OPDA treatment in comparison to DEGs induced by wounding showed a significant induction of a sub-cluster of genes belonging to sulfate reduction and assimilation pathway, the phosphorelay signal transduction pathway as well as the hormone-mediated signaling pathway (Supplementary Fig. 7b, Fig. 4a, Supplementary Dataset 3). These genes were not induced by the endogenous rise of OPDA and included genes encoding sulfate reductases, such as SULFATE-DEFICIENCY INDUCED (SDI1), APS REDUCTASE3 (APR3), and RESPONSE TO LOW SULFUR3 (LSU31), and ORA59, an APETALA2/ETHYLENE RESPONSE FACTOR (AP2/ERF) domain transcription factor which is involved in JA and ethylene signaling and in defense ^46^ (Fig. 4b). This is in line with the previously reported induction of sulfur metabolism by OPDA binding to the CYP20-3 module ^35^. Here, the activation of the sulfur metabolism pathway was a specific effect of exogenously supplied OPDA that did not correlate with an endogenous effect of the compound. Formerly described OPDA-induced genes were induced by both OPDA treatment and wounding of the *opr2opr3* mutant, however, their wound induction in *aos* indicated their involvement in general stress-response pathways (Fig. 2, Fig. 4d). In addition, OPDA application to *opr2opr3* seedlings resulted in a slight, but significant induction of JA signaling genes (Supplementary Fig. 7a, b, Supplementary Dataset 3). To get a deeper insight into this JA-mediated response occurring in *opr2opr3* seedlings upon OPDA treatment, expression levels of JA responsive genes were determined in comparison to wounded seedlings from wild-type and both mutants (Supplementary Fig. 7c). OPDA-treated *opr2opr3* and wounded wild-type seedlings shared the up-regulation of several JA marker genes, such as those encoding JAZ transcription factors and JA biosynthesis enzymes (Supplementary Fig. 7c). Validation of transcript accumulation of *JAZ2*, *JAZ7*, *JAZ13* and *CHL1* by RT-qPCR confirmed their exclusive induction in *opr2opr3* seedlings upon OPDA application but not upon endogenous OPDA accumulation by wounding (Supplemental Fig. 7d). This hints towards the conversion of the exogenously applied but not the endogenously formed OPDA to JA/JA-Ile despite the loss of OPR3 and OPR2 in the *opr2opr3* background. OPDA and JA levels in *opr2opr3* seedlings determined after wounding and OPDA treatment showed unequivocally that *opr2opr3* seedlings accumulated slowly higher levels of JA-Ile following OPDA application in comparison to wounding (Supplemental Fig. 7e). Despite these levels being lower than the JA-Ile levels in wounded wild-type seedlings, they correlated with the activated JA signaling in *opr2opr3* seedlings after OPDA application. Interestingly, the OPDA-fed *opr2opr3* seedlings accumulated only slightly higher OPDA levels than the wounded seedlings, whereas *dn*-OPDA accumulated significantly only following endogenous OPDA formation after wounding (Supplemental Fig. 7e). This suggests that the exogenously applied OPDA is either detoxified by conjugation to GSH ^47^ or rapidly converted to 4,5-ddh-JA as shown by Ueda et al. (accompanying manuscript). In contrast, *dn*-OPDA accumulating after wounding in *opr2opr3* seedlings might be produced from OPDA entering the β-oxidation cycle ^43^ or from the parallel hexadecanoid pathway ^23^. These results raise the question whether the different patterns of gene expression detected here, are due to a compartmentalization of OPDA synthesized following wounding, thereby not inducing the responses that the exogenously supplied OPDA does.

**Fig. 4:**
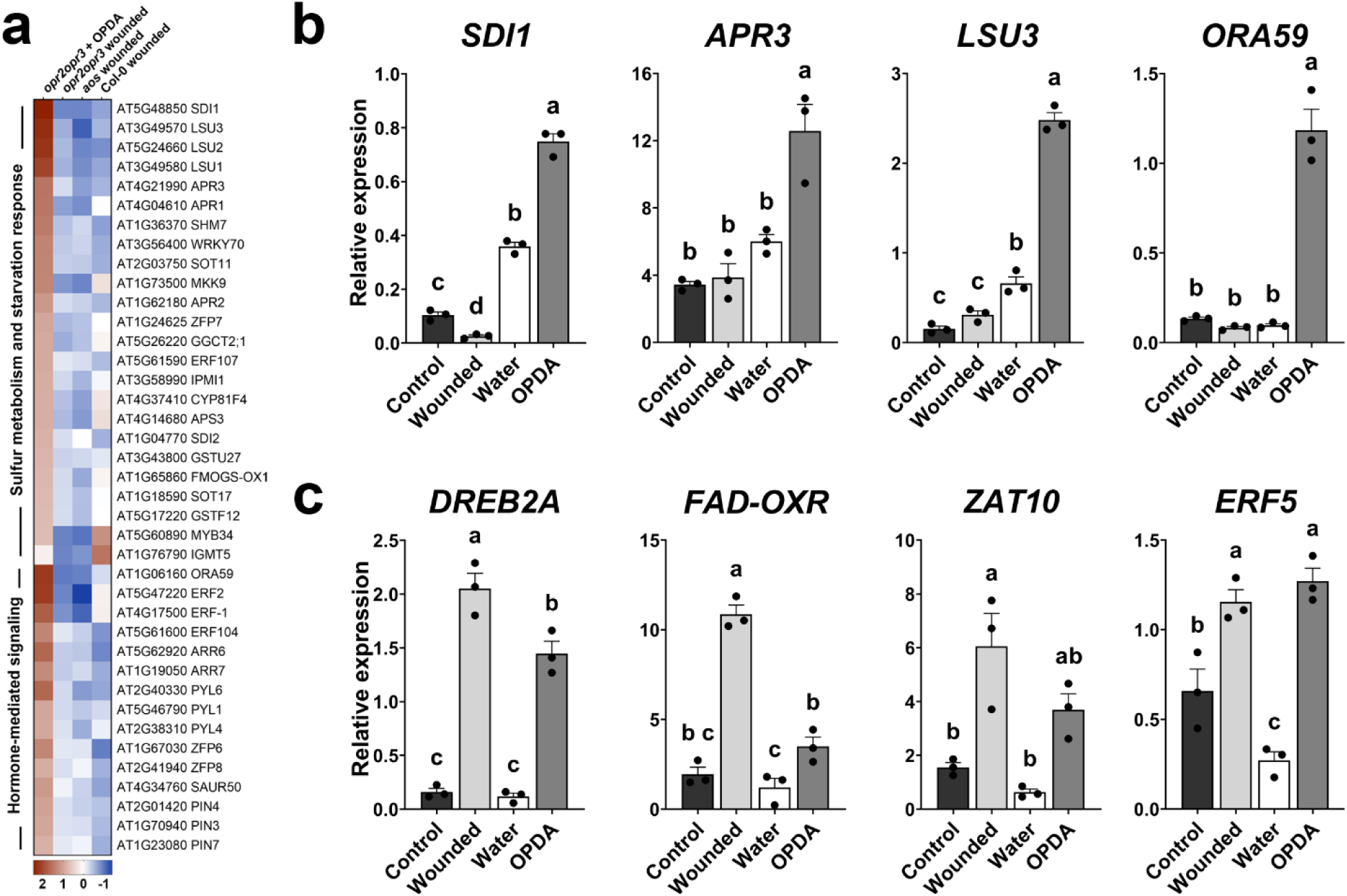
Application of OPDA results in the induction of sulfur metabolism. **(a)** Heatmap of DEGs showing enrichment in genes involved in sulfur assimilation and common hormone signaling in seedlings of *opr2opr3* following application of 25 µM OPDA for 30 min. DEGs were plotted and compared to those from wounded seedlings of Col-0, *opr2opr3* and *aos* using average FPKM values transformed into row-normalized Z-score of three biological replicates. **(b)** RT-qPCR validation of the up-regulation of the sulfur metabolism genes *SDI1*, *APR3* and *LSU3* as well as the regulator of JA- and ethylene-responsive gene expression *ORA59* in *opr2opr3* seedlings treated with OPDA in comparison to wounded *opr2opr3* seedlings. OPDA-treatment and wounding of seedlings were compared to seedlings which were non-wounded (control) and water-treated (water), respectively. **(c)** RT-qPCR validation of the up-regulation of the OPDA-responsive genes *DREB2A*, *FAD-OXR*, *ZAT10* and *ERF5* in *opr2opr3* seedlings treated with OPDA in comparison to wounded *opr2opr3* seedlings. OPDA-treated and wounded *opr2opr3* seedlings were compared to their respective controls, non-wounded (control) and water-treated seedlings, respectively. Bars **(b-c)** represent means of three biological replicates with 120 seedlings each (single dots; ±SEM). Statistically significant differences among treatments were calculated using Two-Way ANOVA followed by Tukey-HSD and are indicated by different letters.

### Test for subcellular distribution of the endogenously produced OPDA using a trans-organellar complementation of the *opr2opr3* mutant

The data obtained from wounded seedlings of *opr2opr3* and described above suggest that endogenously produced OPDA does not exhibit signaling capacity. This leads to the question, whether OPDA is restricted to the cell compartments of its synthesis, transfer and metabolization, namely plastids, cytosol and peroxisomes, respectively. To test its putative translocation to other cell compartments, a trans-organellar complementation approach according to ^48^ using the *opr2opr3* mutant was performed. OPR3 was targeted to different organelles in the *opr2opr3* mutant, and the fertility of these plants was checked. A rescue of the JA-deficiency phenotype of the *opr2opr3* mutant would indicate whether its substrate, OPDA, is able to translocate to the tested organelles (Supplementary Fig. 8).

OPR3 is a peroxisome-localized protein and is imported into the matrix of peroxisomes through the peroxisomal targeting signal type 1 (PTS1) consisting of a tripeptide at the extreme carboxy-tail of the protein ^49^. OPR3 contains a C-terminal SRL, which is a variation of the prototypic PTS1 signal SKL. Removal of this signal resulted in cytosolic localization of a fusion of OPR3ΔSRL with YFP when transiently expressed in *Nicotiana benthamiana* leaves, indicating that import into peroxisomes was abolished (Fig. 5a). Targeting of OPR3 to the nucleus, cytosol, plastids, mitochondria, and endoplasmic reticulum (ER) was done by fusing OPR3ΔSRL with the respective targeting signals or target peptides (Supplementary Tab. 5). Validation of the subcellular localization of the different organelle variants of OPR3 with co-localization studies in *N. benthamiana* protoplasts was performed using established organelle markers ^50^ and indicated correct subcellular localizations of OPR3 compared to the markers independently of the position of the YFP fluorescent tag (Supplementary Fig. 9). Stable transgenic *opr2opr3* lines complemented with the organelle variants of OPR3ΔSRL fused to YFP were generated and resulted in clear subcellular localizations of OPR3-YFP to the nucleus, cytosol, chloroplast stroma, mitochondria, and ER (Fig. 5b). As tested by RT-qPCR, all transgenic lines exhibited a similar level of transgene expression (Supplementary Fig. 10).

**Fig. 5:**
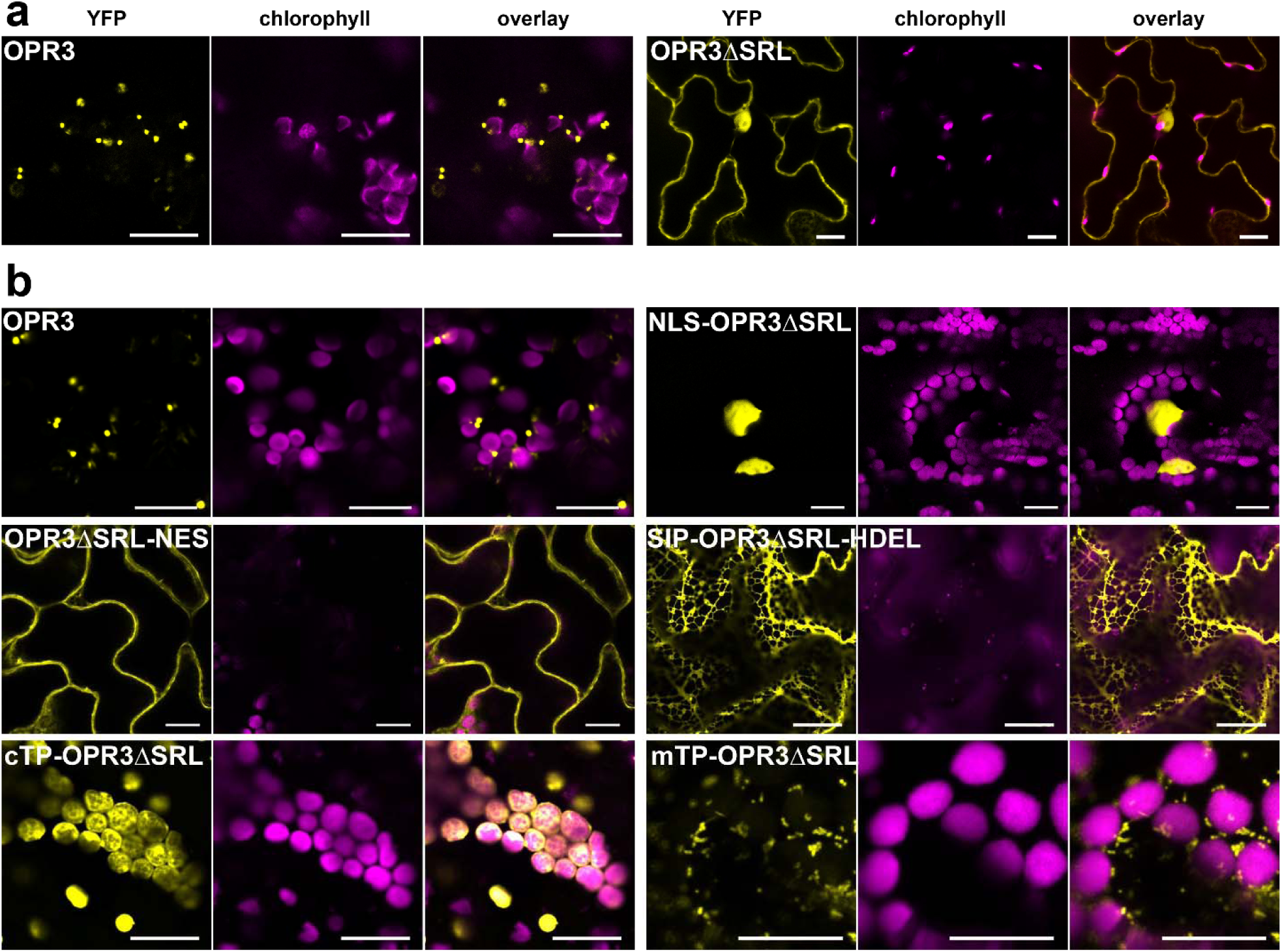
The removal of the SRL peroxisomal signal results in a cytosolic localization of OPR3 in *N. benthamiana* leaves and targeting of OPR3ΔSRL to the nucleus, cytosol, plastid stroma and mitochondria in stable transformed *opr2opr3* lines. **(a-b)** OPR3 **(a)** and OPR3ΔSRL **(b)** were C-terminally fused to YFP and transiently expressed in *N. benthamiana* leaves under control of the CaMV *35S* promoter. Imaging was performed at 48-72 h post infiltration. **(c-g)** Stable *opr2opr3* complementation lines expressing OPR3ΔSRL targeted to the nucleus **(c)**, the cytosol **(d)**, ER **(e)**, chloroplast stroma **(f)** and mitochondria **(g)**, all under control of the CaMV *35S* promoter. All pictures are taken from homozygous T2 single insertion lines. The yellow signal corresponds to YFP, whereas the magenta corresponds to chlorophyll autofluorescence. Scale bars represent 20 µm. NLS, nuclear targeting signal; NES, nuclear export signal; SIP, signal peptide; cTP, chloroplast targeting peptide; mTP, mitochondria targeting peptide.

The main phenotypic feature of JA-deficient and JA-insensitive Arabidopsis mutants is the male sterility, which is also characteristic for *opr2opr3* plants and is visible by defects in filament elongation, pollen release and seed set ^41^. Plants of *opr2opr3* transformed with an empty vector (EV) showed short stamens with non-dehiscing anthers (Fig. 6a, b), similar to the reported male sterility phenotype of the *opr3* mutant ^38^. As a positive control, the complementation of *opr2opr3* with the peroxisome targeted OPR3 showed full rescue of the flower phenotype (Fig. 6c). Interestingly, all other tested organelle-variants of OPR3 also rescued the flower sterility and showed open flowers like the wild-type with elongated stamens and anthers releasing pollen (Fig. 6d-h). This rescue of sterility translated also in the ability of the lines to form siliques containing seeds. The only exception were plants complemented with the ER-targeted OPR3 having partial rescue of flower fertility only. These plants formed fewer and smaller siliques compared to the wild type (Fig. 6f). This suggests that OPR3 localized to the ER may have restricted access to its substrate, potentially limiting JA/JA-Ile biosynthesis compared to other subcellular compartments.

**Fig. 6:**
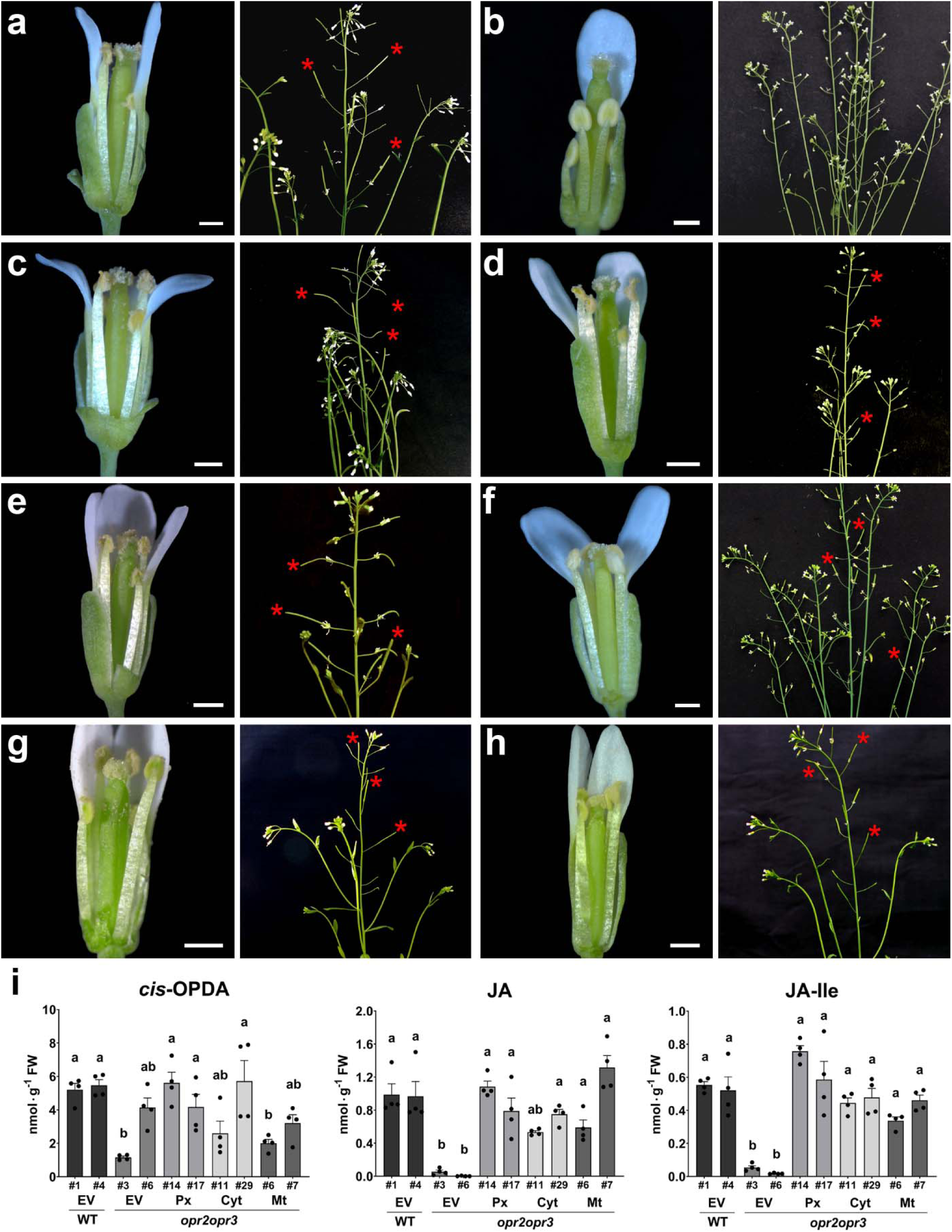
Restitution of the fertility and JA/JA-Ile levels of *opr2opr3* plants by OPR3-targeted to different organelles. **(a-h)** Open flower buds showing stamen filament elongation and pollen release by anthers (scale bars represent 0.5 mm) and plant inflorescences showing silique formation at days 50-60 post germination with asterisks marking fully developed siliques. Col-0 and *opr2opr3* transformed with an empty vector **(a, b)** show fertility and sterility, respectively. *opr2opr3* T2 lines complemented with peroxisome- **(c)**, nucleus- **(d)**, cytosol- **(e)**, ER- **(f)**, plastid stroma- **(g)** and mitochondria- **(h**) targeted OPR3 fused to YFP show the rescue of the *opr2opr3* sterility phenotype. Red asterisk show exemplary siliques containing seeds. **(i)** OPDA, JA and JA-Ile accumulation at 1 h after wounding of seedlings of wild-type (WT) or *opr2opr3* transformed with either empty vector (EV) or *35S::OPR3* targeted to peroxisomes (Px), cytosol (Cyt) or mitochondria (Mt). Numbers refer to independent transgenic lines, which were selected for single insertion and homozygosity. Bars represent means of four biological replicates with 120 seedlings each (single dots; ±SEM). Statistically significant differences between *opr2opr3* transformed with empty vector and transformed with the OPR3 constructs were calculated using One-Way ANOVA followed by Tukey HSD (p<0.05) and are denoted by different letters.

Reconstitution of the JA pathway in vegetative tissue of the *opr2opr3* complementation lines was analyzed by determination of wound-induced levels of OPDA, JA and JA-Ile in two-week-old seedlings. Hormone levels of *opr2opr3* complemented with OPR3 targeted to either peroxisomes, cytosol or mitochondria were compared to those of wild-type and *opr2opr3* transformed with an empty vector. Data from two independent transformed lines each revealed that complementation of *opr2opr3* with OPR3 restored the JA/JA-Ile levels to wild-type levels (Fig. 6i). These results show the reconstitution of JA biosynthesis by OPR3 localized to all the tested compartments.

### Trans-organellar complementation with OPR2 and OPR1 to determine whether OPDA or 4, 5-ddh-JA is the translocating substrate

JA biosynthesis in *opr2opr3* complemented with OPR3 located in different organelles would occur only if its substrate, OPDA, is able to translocate to these compartments. However, as an alternative pathway, OPDA could enter directly the β-oxidation as reported recently ^43^. This would result in formation of 4,5-ddh-JA, which might be metabolized by OPR3 irrespective of the organellar location. Conversion of 4,5-ddh-JA to JA by OPR3 was tested by *in vitro* enzymatic assay and showed that OPR3 reduces 4,5-ddh-JA *in vitro*, however to a lesser extent than OPDA (Supplementary Fig. 11). Based on substrate consumption after 10 min, only half of the given 4,5-ddh-JA was reduced while OPDA was completely converted to 3-oxo-2-(20(Z)-pentenyl)-cyclopentane-1-octanoic (OPC-8). The control reactions with inactivated enzyme showed no reduction activity. The results clearly show that OPR3 is able to reduce both cyclopentenones, 4,5-ddh-JA and OPDA, *in vitro*. In addition, it has been shown by recombinant enzymatic assays that OPR2 but not OPR1 is able to reduce 4,5-ddh-JA ^43^. Conversely, OPR1 is able to reduce OPDA although with very low efficiency but OPR2 is not ^21^. To determine whether OPDA or 4,5-ddh-JA is the substrate able to translocate to the tested cell compartments and specifically to cell compartments unrelated to the location of JA biosynthesis, peroxisomal, cytosolic and mitochondrial variants of OPR1 and OPR2, both fused to YFP, were used to complement the *opr2opr3* mutant. Both OPRs localized to the cytosol, whereas addition of a C-terminal SKL and an N-terminal mitochondrial target sequence led to localization in peroxisomes and mitochondria, respectively (Fig. 7a). All three variants expressing OPR2 rescued the flower sterility and showed open flowers like the wild-type with elongated stamens and anthers releasing pollen (Fig. 7b) as well as formation of seed-containing, fully developed siliques (Fig. 7c). In contrast, expression of OPR1 located to peroxisomes was the only OPR1 variant leading to a rescue of the flower fertility, whereas targeting OPR1 to the cytosol led to partial rescue of flower fertility only and targeting OPR1 into the mitochondria did not restore the fertility (Fig. 7b, c). These data were supported by the fact that only seedlings harboring the OPR2 variants and peroxisomal-located OPR1 were able to accumulate JA and JA-Ile upon wounding (Fig. 7d), although all lines accumulated similar levels of OPDA (Supplementary Figure 12). The JA/JA-Ile levels were lower than those of wounded wild-type seedlings, but the significantly increased levels in comparison to *opr2opr3* transformed with the empty vector showed the capability of these plants to produce JA/JA-Ile. Most importantly, however, expression of OPR1 located to mitochondria did not restore fertility nor JA/JA-Ile levels. Since OPR1 does not convert 4,5-ddh-JA to JA, this result points to the fact that 4,5-ddh-JA but not OPDA might be the compound to be converted by OPR2 or OPR3 in different organelles.

**Fig. 7:**
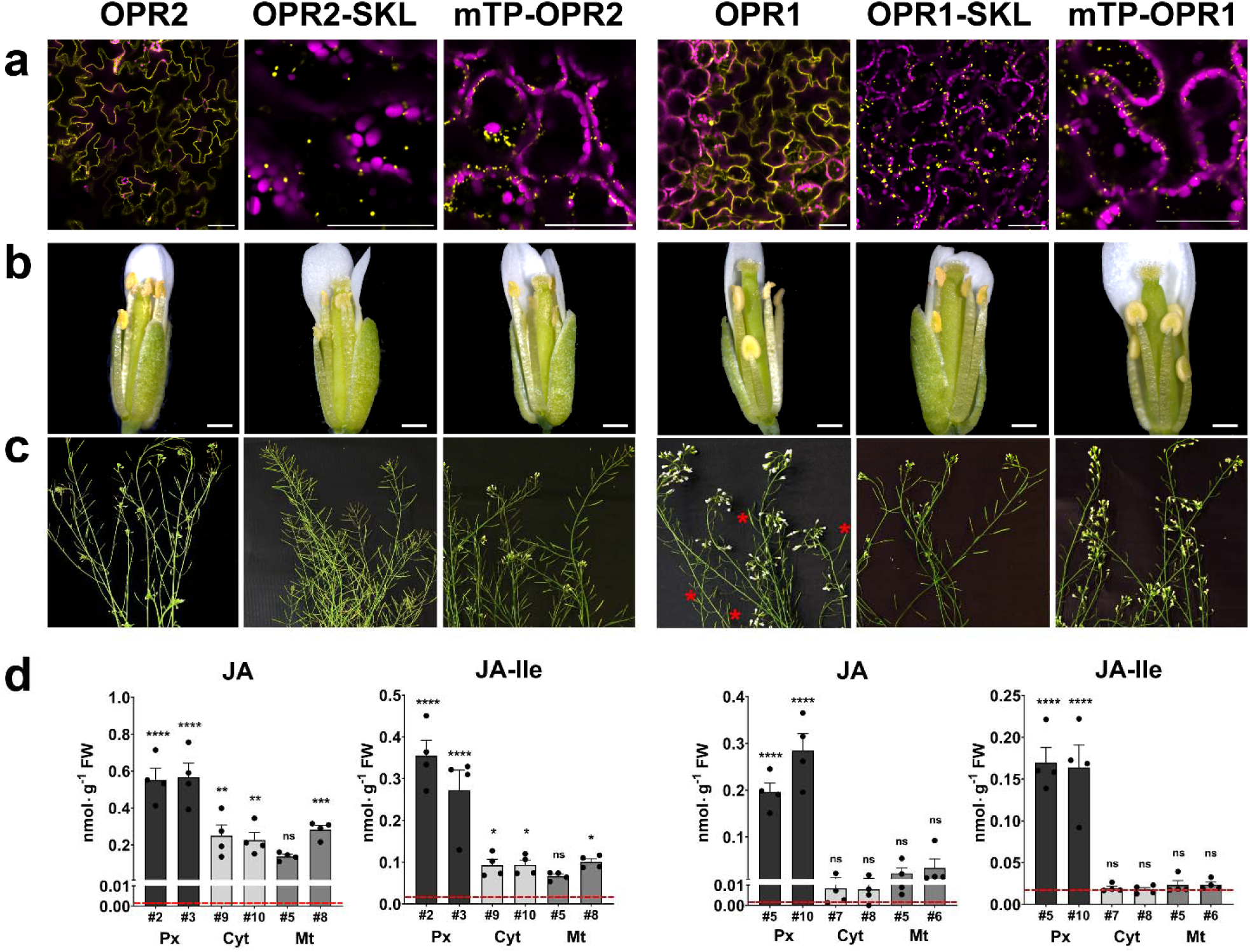
Restitution of the fertility and JA/JA-Ile levels of *opr2opr3* plants by OPR2, but not by OPR1, both targeted to different organelles. Stable *opr2opr3* complementation lines expressing OPR2 (left) or OPR1 (right) targeted to the cytosol (without any signal peptide), to the peroxisomes (OPR-SKL) or to mitochondria (mTP-OPR), all under control of the CaMV *35S* promoter. All transgenic lines showed a similar level of transgene expression (Supplementary Fig. 10). **(a)** Micrographs showing the correct localization of the respective OPR-YFP-fusion. The yellow signal corresponds to OPR-YFP, whereas the magenta signal corresponds to chlorophyll autofluorescence. Scale bars represent 50 µm. **(b, c)** Open flower buds to show stamen filament elongation and pollen release by anthers **(b)** and plant inflorescences showing silique formation at days 50-60 post germination **(c)**. Note that transformation of *opr2opr3* with all variants of OPR2 and with OPR1 targeted to peroxisomes resulted in a rescue of fertility visible by elongated filaments and fully developed siliques. Targeting of OPR1 to cytosol led to partial rescue shown by few fully developed siliques only (red asterisks), whereas targeting of OPR1 to mitochondria did not rescue fertility. Scale bars in **(b)** represent 0.5 mm. **(d)** JA and JA-Ile accumulation at 1 h after wounding of seedlings of *opr2opr3* transformed with either *35S::OPR2* or *35S::OPR1*, either targeted to cytosol (Cyt), peroxisomes (Px), or mitochondria (Mt). Numbers refer to independent transgenic lines, which were selected for single insertion and homozygosity. The mean levels of JA/JA-Ile of *opr2opr3* transformed with the empty vector are taken from Fig. 6 and are represented by red dashed lines for comparison. Bars represent means of four biological replicates with 120 seedlings each (single dots; ±SEM). Statistically significant differences between *opr2opr3* transformed with the different *OPR2* and *OPR1* constructs and *opr2opr3* transformed with the empty vector were calculated using One-Way ANOVA with Fisher LSD post-hoc test and are indicated by asterisks with *p<0.05, **p<0.01, ***p<0.001, and ****p<0.0001; ns = non-significant.

## Discussion

JA plays a major role in growth and defense against pathogens, herbivory attacks and mechanical wounding. Its signaling through the bioactive JA-Ile is well established in Angiosperms, while in non-vascular plants, such as *M. polymorpha*, OPDA, *dn*-*iso*-OPDA and Δ^4^-*dn*-*iso*-OPDA are bioactive^13,24–26,51^. Given that *A. thaliana* plants produce OPDA and *dn*-OPDA, possible function of OPDA as a signaling molecule independently from JA was proposed. OPDA was shown to induce a distinct transcriptional change from JA when exogenously applied to Arabidopsis plants ^32,45^. Additionally, OPDÁs involvement in processes like seed germination and stomatal closure was demonstrated ^29–31^. The mechanism by which OPDA exerts these functions in vascular plants is, however, still elusive ^17^. A major genetic background used for studying OPDA functions was the *opr3* mutant ^41^. Recently, an OPR3-independent pathway of JA formation through the cytosolic isoenzyme OPR2 has been identified ^43^. Hence, uncoupling of JA production from OPDA was not achieved in the *opr3* mutant, and the characterized functions of OPDA cannot be attributed to OPDA with certainty. Here, we addressed the question of whether OPDA functions independently from JA as a signaling molecule in Arabidopsis by utilizing the *opr2opr3* mutant, which does not produce JA/JA-Ile upon wounding ^43^.

The use of the *opr2opr3* mutant is limited by its lower production of OPDA in comparison to the wild type due to the disrupted JA positive feedback loop as indicated by reduced protein levels of the biosynthesis enzymes, such as AOC ^1^. The enhancement of the AOC levels and subsequent OPDA production in the JA-deficient mutants was achieved by restituting the JA feedback loop through a repetitive supply of MeJA to seedlings during development (Fig. 1). This result validated the role of the JA positive feedback loop in the regulation of the basal levels of the biosynthetic enzymes and setting the mutant ability to produce OPDA to similar levels as wild-type plants do ^44^.

### The wounding response of wild-type seedlings involves JA-dependent and independent processes

To inspect the separate roles of OPDA and JA upon wounding, a comparative analysis of the transcriptomes of seedlings of wild type, *opr2opr3* and *aos* was carried out. The transcriptomic change induced by wounding in wild-type seedlings involved an up-regulation of genes previously identified as JA-dependent or associated with immune responses, validating the specific signaling role of JA-Ile in the wound response of seedlings as it has been described for mature leaves of Arabidopsis and other plant species ^5,52–54^. Unexpectedly, seedlings of the *aos* mutant being deficient in synthesis of JA and OPDA, also showed a major transcriptional change upon wounding that was common to the *opr2opr3* mutant and the wild type (Fig. 2). The transcriptional response was significantly enriched in general stress response pathways, with an induction of oxidative stress- and transcription-related genes. This implies that JA-independent signaling processes occur in response to mechanical wounding leading to transcriptional reprogramming. Among such processes, water stress might contribute to the regulation of wound-responsive genes in a JA-independent manner ^55^. Ethylene and abscisic acid (ABA) also contribute to the wound response by regulating photosynthesis and drought responsive genes, respectively ^56^. An ABA-dependent and JA-independent transcriptional regulation of wax biosynthesis to seal the wounded sites of Arabidopsis leaves has been recently characterized, but appears to be controlled post-translationally by JA ^57^. As plants defective in JA/JA-Ile biosynthesis or perception are severely diminished in their defense response ^41,58–60^, the JA-independent wound response has rather limited contribution to plant defense and might predominantly mitigate the adverse effects of wounding itself by promoting wound healing and tissue regeneration ^61^.

### An OPDA-specific transcriptional response is missing in wounded *opr2opr3* seedlings, but specific pathways are activated by the exogenous application of OPDA

The general stress response that was both JA and OPDA-independent, comprised genes that were previously identified as OPDA-specific response genes, such as *DREB2A*, *FAD-OXR*, *ZAT10, ERF5, TCH4,* and *GST6* ^32^. These genes did not show differential expression when the transcriptomes of *opr2opr3* and *aos* were compared indicating that their induction by wounding is OPDA-independent (Fig. 2). Moreover, these genes were induced by wounding also in leaves of two *coi1*-mutant lines excluding any induction by JA/JA-Ile possibly caused by an insufficient JA-deficiency of the mutants. Several abiotic stresses were reported to up-regulate the same genes ^62–65^, further indicating their independence from OPDA. Moreover, direct comparison of transcriptomes from wounded *opr2opr3* and *aos* seedlings with standard cut-off parameters did not yield a significant list of putative OPDA-regulated genes further confirming the absence of an OPDA signaling (Fig. 3). This could be due to three facts: (i) A putative receptor for OPDA might be missing in *A. thaliana*, (ii) OPDA is converted to 4,5-ddh-JA, which does not accumulate to levels having signaling capacity ^43^, and (iii) OPDA remains largely sequestered in the plastids by its biosynthetic enzymes ^66^. Regarding perception of OPDA, evolutionary studies of the JA-Ile co-receptor COI1 showed that the Arabidopsis protein does not perceive OPDA or *dn*-OPDA, dismissing possible signaling of OPDA through COI1 in this plant ^13^. Moreover, it has been shown that *dn*-*iso*-OPDA rather than *dn*-OPDA is the bioactive molecule in mosses, and that this compound does not occur in *A. thaliana* ^26^. The transcriptomic data obtained from *opr2opr3*, *aos*, *coi1-16* and *coi1-30* therefore point to the fact that OPDA or its homolog *dn*-OPDA are not perceived by COI1 or any other receptor in Arabidopsis.

The transcriptional signature of OPDA was predominantly observed in experiments relying on its exogenous application ^32,35,45^. Therefore, we compared the transcriptional changes following application of OPDA to *opr2opr3* seedlings with those by wound-induced, endogenously produced OPDA. This comparison revealed an up-regulation of sulfur assimilation and GSH production by OPDA being attributed as specific signaling functions of OPDA ^35^. Here, the transcriptional activation of sulfur metabolism pathway was exclusive to the exogenous OPDA supply, contradicting an intrinsic signaling function of the endogenous OPDA (Fig. 4). As an electrophilic species, OPDA was shown to disturb redox homeostasis ^67^ and to inhibit photosynthesis by affecting the photosystem II fluorescence ^33,68^. In parallel, the *in planta* conjugation of OPDA to GSH, which is an important antioxidant substance, and its degradation in the vacuole were also demonstrated ^47^. This implies that OPDA may not directly regulate the sulfur assimilation pathway but instead triggers this response through its potential detoxification when exogenously applied to plants, suggesting that the observed transcriptional response to exogenously applied OPDA could be a side effect rather than a direct effect of OPDA itself. The main route for the conversion of applied OPDA seems to be, however, its direct conversion to 4,5-ddh-JA, which then reaches levels up to 4000 pmol per g FW (Ueda et al., accompanying manuscript). Moreover, Ueda et al. showed convincingly that OPDA applied to Arabidopsis seedlings does not serve as genuine bioactive form responsible for the expression of OPDA-marker genes. As shown by Ueda et al., 4,5-ddh-JA is inducing the expression of OPDA-specific response genes in a dose-dependent manner, possibly due to its electrophilic properties. Upon wounding of *opr2opr3*, however, 4,5-ddh-JA accumulates up to 500 pmol per g FW ^43^, a level that does not seem to be sufficient to induce a specific gene expression (Ueda et al., accompanying manuscript). Here, the general stress response might be the main factor for transcriptional changes including the previously identified OPDA-marker genes as depicted from the wound response of *aos* mutant plants. These genes, being responsive to several abiotic stresses ^62–64^, including wounding in a JA-independent manner (as shown in this manuscript), further suggest their unspecific response to OPDA. This rather indicates that applying electrophilic compounds, such as 4,5-ddh-JA or OPDA, causes oxidative stress, resulting in the induction of general stress-responsive genes. Next to jasmonates, there are other compounds known that mediate at least a part of the wound response, such as reactive oxygen species (ROS) ^69^ and γ-aminobutyric acid (GABA) ^70^. These JA-independent responses might be necessary for wound-healing, which seems to be a process independent on the JA-dependent defense response ^71^.

### Endogenously formed OPDA is sequestered to the organelles of JA biosynthesis

To address the question whether wound-induced OPDA remains largely sequestered in the plastids or whether it is converted to 4,5-ddh-JA similarly to the applied OPDA (Ueda et al., accompanying manuscript), a trans-organellar complementation approach was performed by directing OPR3 into various organelles (Figs. 5, 6, Supplementary Fig. 9). Occurrence of OPR3 in any of the tested organelles resulted in a complementation of the phenotype and JA/JA-Ile production upon wounding in *opr2opr3* mutants. This led to the assumption that OPDA might travel throughout the cell. Nevertheless, the possibility still exists that OPDA is translocated rapidly via the cytosol to the peroxisomes, where it is converted to 4,5-ddh-JA by β-oxidation as shown previously for *opr3* and *opr2opr3* ^43^. Given that OPR3 is able to convert 4,5-ddh-JA to JA and OPDA to OPC-8 *in vitro* (Supplementary Fig. 10), the complementation approach with OPR3 could not discriminate between the presence of either OPDA or 4,5-ddh-JA outside of the plastids or the peroxisomes. To address this, organelle variants of the isoenzymes OPR2 and OPR1 were used to complement the *opr2opr3* mutant (Fig. 6). OPR2 and OPR1 differ in their enzymatic activity against OPDA and 4,5-ddh-JA: OPR2 is able to convert both compounds, but OPR1 has been shown to be inactive on 4,5-ddh-JA ^43^. Since OPDA and 4,5-ddh-JA should occur in peroxisomes and in the cytosol, reconstitution of JA biosynthesis was obtained using either OPR1 or OPR2 targeted to peroxisomes or the cytosol. Targeting OPR1 or OPR2 to mitochondria, however, discriminated whether OPDA or 4,5-ddh-JA is the substrate able to translocate to other organelles (Fig. 8). Here, complementation of *opr2opr3* with a mitochondrial-located OPR1 did not rescue fertility and JA/JA-Ile production (Fig. 7), whereas OPR2 in mitochondria did. Additionally, OPR1 fully complemented the mutant only within the peroxisome, and its cytosolic variant resulted in minimal JA-Ile formation and partial rescue of fertility. These results suggest a limited occurrence of endogenously formed OPDA in the cytosol and its preferred compartmentalization in plastids and peroxisomes, whereas 4,5-ddh-JA might be the substrate able to translocate between cell organelles. The possible translocation of 4,5-ddh-JA correlates with its potential role in signaling or its effect as an electrophilic compound, as demonstrated by Ueda et al. (accompanying manuscript).

**Fig. 8:**
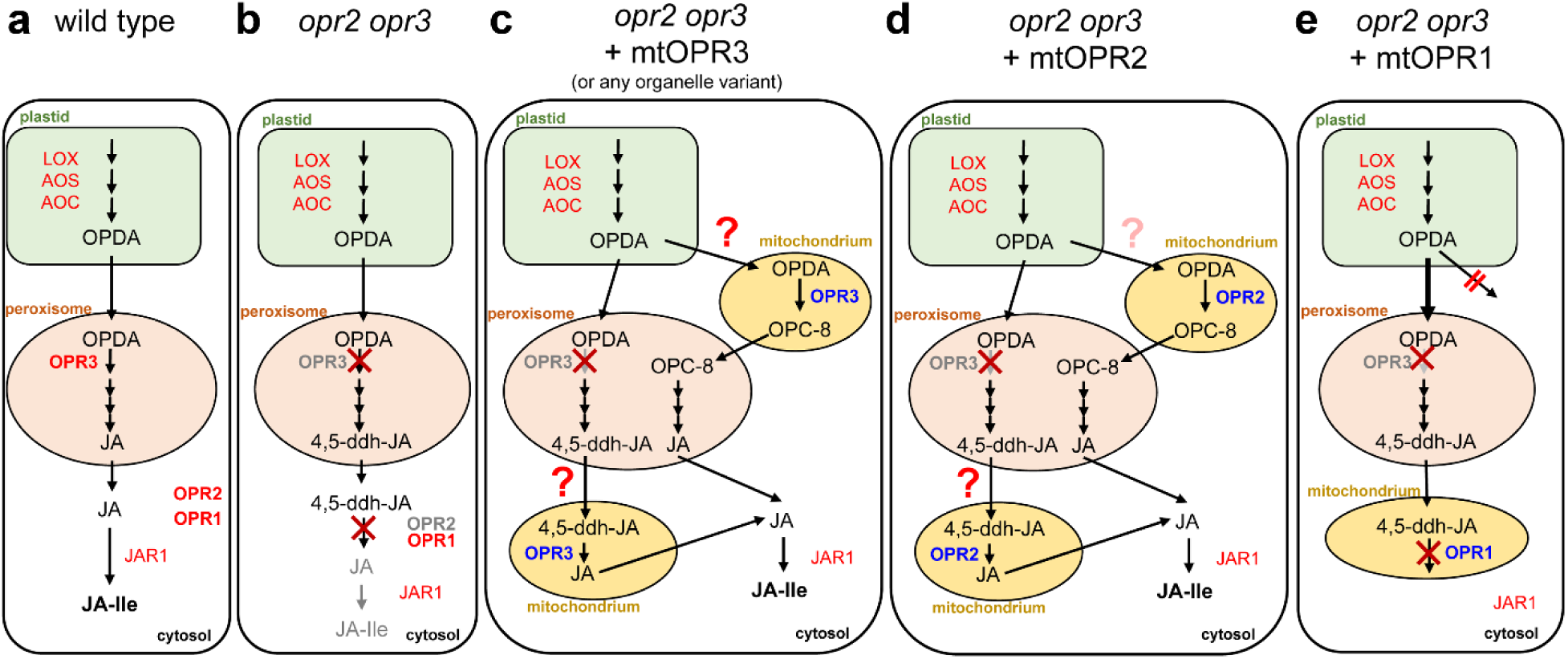
Schematic picture on the approaches leading to insights about the putative distribution of OPDA and 4,5-ddh-JA within cells of a wounded Arabidopsis seedling. Putative subcellular distribution of OPDA and its conversion to JA deduced from the trans-organellar complementation approach. **(a)** In wild-type plants, OPDA is converted to JA in the peroxisome through the OPR3 canonical route, the contribution of OPR2 located in the cytosol appears to be neglectable but comes into place if OPR3 is not functional (e.g., *opr3* single mutant, see ^42^). **(b)** The interruption of both routes in the *opr2opr3* mutant results in conversion of OPDA to 4,5-ddh-JA which is not metabolized to JA/JA-Ile. **(c)** Targeting of OPR3 to any organelle in *opr2opr3* mutant plants (depicted for mitochondrial localization) restitutes JA biosynthesis in this mutant. If this is a result of either an ability of OPDA or of its β-oxidation product, 4,5-ddh-JA, to translocate to other cell compartments cannot be decided by this approach, since OPR3 is active to convert OPDA and 4,5-ddh-JA (red question marks). Trans-organellar complementation by using OPR2 or OPR1 targeted into mitochondria, however, helped to clarify this point. **(d)** Mitochondrial targeting of OPR2 with its specificity to 4,5-ddh-JA complemented JA biosynthesis favoring rather an ability of 4,5-ddh-JA to be mobile, although the conversion of OPDA can still not be excluded due to some activity of OPR2 towards OPDA (light red question mark). **(e)** As final proof, OPR1 was targeted to mitochondria, because OPR1 is specific for OPDA and does not convert 4,5-ddh-JA. Expression of mtOPR1 in *opr2opr3* did not restitute JA/JA-Ile production. Even the very limited JA formation by cytosolic OPR1 suggests restricted occurrence of OPDA in this compartment (not shown in this scheme, data see Figure 7). The lack of JA formation by mitochondrial OPR1 suggests that OPDA may not have the ability to access this cellular compartment and might largely compartmentalized in the organelles involved in JA biosynthesis, the plastids and peroxisomes, and occurs in limited amounts in the cytosol.

In conclusion, despite its perception and signaling in non-vascular plants, OPDA itself does not mediate transcriptional changes in Arabidopsis. The induction of a specific transcriptional change occurred exclusively upon exogenous supply but not upon endogenous rise of OPDA, although in both cases OPDA is (partially) converted to 4,5-ddh-JA (^43^ and Ueda et al., accompanying manuscript). Here, the endogenous levels of 4,5-ddh-JA – being low upon wounding, but high upon exogenously applied OPDA – might be determining, whether it acts as an electrophilic (toxic) compound inducing expression of stress-related genes. In turn, the data obtained after wounding suggest that the endogenous OPDA is compartmentalized and thereby its level in the cytosol highly regulated. The discrepancies between the effects of endogenously produced and exogenously supplied OPDA suggest an absence of signaling per se and/or a tightly regulated compartmentalization to prevent its effects as an electrophilic species. Both scenarios imply that OPDA does not inherently function as a signaling compound in Arabidopsis, unless such functionality manifests under specific, yet uncharacterized conditions. It is plausible that OPDA-oxidation by JASMONTE-INDUCED DIOXYGENASE1 (JID1) ^72^, or conjugation to amino acids ^73,74^ or GSH ^37^ contribute to the regulation of the cytosolic OPDA level.

## Material and methods

### Plant material, growth conditions, and treatment

*Arabidopsis thaliana* wild-type (ecotype Col-0) and the mutant lines *opr2opr3* ^43^, *aos* (*dde2-2* ^40^, *coi1-16* ^75^ and *coi1-30* ^76^ were genotyped using the primers listed (Supplementary Tab. 6). After surface sterilization with 4% bleach and three-day stratification at 4°C, seedlings were grown for ten days in liquid Murashige and Skoog (MS) medium (pH 5.7, 1% sucrose), whereas adult plants were grown for four weeks in individual pots containing steam-sterilized clay, coir fiber, and vermiculite. Plant growth took place in Phytocabinets (Percival Scientific, www.percival-scientific.com/) at 120 µE m^-2^ s^-1^ under short day conditions (10/14 hours light/dark cycle) at 21/19 °C and 65% relative humidity. For wounding, seedlings were squeezed with forceps with serrated teeth. Phytohormone treatments were done by adding either 25 µM OPDA (≥ 95%, Cayman Chemical, www.caymanchem.com) or 1 µM MeJA to the medium.

### RNA isolation, RNA-seq and quantitative RT-PCR analysis

Total RNA was isolated from homogenized frozen material using the RNeasy Plant Mini Kit (Qiagen) and treated with the DNA-free™ DNA Removal Kit (Invitrogen, #AM1906). First strand cDNA synthesis from 1 µg DNA-free RNA was carried out using the RevertAid H Minus reverse transcriptase with oligo(dT)18 primers (ThermoFisher Scientific™, www.thermofisher.com). Quantitative PCR was carried-out in Hard-Shell® 96-Well PCR Plates (Bio-Rad Laboratories, www.bio-rad.com, #HSP9601) supplied with 10 µL reaction mix of 1.5 ng/µL cDNA, 1x EvaGreen QPCR Mix II (Bio&Sell, www.bio-sell.de, #BS76.580.0200) and 0.2 µM of forward and reverse primers (Supplementary Tab. 6). The reactions were run on a CFX Connect Real-Time PCR Detection System (Bio-Rad Laboratories) with denaturation (95°C for 15 min), amplification (40 cycles of 95°C for 15 s and 60°C for 30 s) and melt curve analysis (95°C for 10 s, 65°C heating up to 95°C with a heating rate of 0.05°C s^−1^). Gene expression was normalized to the housekeeping gene *PROTEIN PHOSPHATASE 2A SUBUNIT A3* (AT1G13320) ^77,78^ using the 2^-ΔCT^ method ^79^ and included biological triplicates.

### Transcriptome analysis

RNA quality and integrity were assessed on Agilent 2100 Bioanalyzer system (Agilent Technologies, www.agilent.com) using the RNA 6000 Nano Kit for standard RNA sensitivity (Agilent, #5067-1511). Three biological replicates were submitted to Novogene (www.novogene.com) for mRNA paired-end short-read sequencing (150 bp length) on an Illumina NovaSeq 6000 Sequencing System and bioinformatics analysis according to their pipeline. Reads were mapped to the *A. thaliana* TAIR10 reference genome. Gene expression levels were determined using the FPKM (Fragments per kilobase per million) method. Gene expression heatmaps, principal component analysis (PCA), gene clustering, and Gene Ontology (GO) enrichment analysis were generated using the iDEP.96 (http://bioinformatics.sdstate.edu/idep96/), iDEP 1.1 (http://bioinformatics.sdstate.edu/idep11/) and ShinyGO 0.80 (http://bioinformatics.sdstate.edu/go/) software tools ^80,81^.

### Protein extraction and immunoblotting

Proteins were extracted by incubation of 50 mg frozen material in 200 µl of extraction buffer (25 mM Tris-Cl pH 6.8, 1% SDS, 1% [v/v] β-mercaptoethanol) at 95°C for ten minutes. Extracts were treated with 1% (v/v) Halt™ Protease and Phosphatase Inhibitor Cocktail (ThermoFisher Scientific™, #78440) and quantified at 595 nm on the SPARK® multimode microplate reader (TECAN) using the Pierce™ Bradford Plus Protein Assay Kit (ThermoFisher Scientific™, #23236). Ten micrograms of total protein were incubated at 96°C for 10 min in Laemmli Buffer (1:1, ^82^ to dissolve AOC trimers ^83^. Proteins were separated on SDS–PAGE (4% and 12% acrylamide for stacking and resolving gels, respectively), transferred to a PVDF membrane and detected using Ponceau S staining or No-Stain™ Protein Labeling Reagent (ThermoFisher Scientific™, #A44449). Membrane blocking in 5% (w/v) BSA in TBST (20 mM Tris–Cl pH 7.8, 150 mM NaCl, 0.05% [v/v] Tween) was followed by immuno-staining using anti-AtAOC primary antibody (1:5000, ^44^ and a goat anti-rabbit IgG secondary antibody conjugated with alkaline phosphatase (1:4000, Chemicon®, Sigma-Aldrich, www.sigmaaldrich.com, #AP307P). Detection by chemiluminescence with Immun-Star AP Substrate (ThermoFisher Scientific, #1705018) was visualized using a Fusion FX Imaging system (Vilber, www.vilber.com). AOC protein bands were quantified relative to total protein loadings using densitometry analysis with ImageJ (https://imagej.nih.gov/ij/index.html).

### Cloning of the OPRs and plant transformation

The OPR3 (1176 bp), OPR2 (1125 bp), and OPR1 (1194 bp) coding sequences (CDS) without stop codons were PCR-amplified from Col-0 cDNA using the Q5® High-Fidelity DNA Polymerase (Bio Labs, www.biolabs.io, #M0491) and the listed primer pairs (Supplementary Tab. 7). For OPR3ΔSRL, the last nine nucleotides were removed, and silent mutations C205G and G1109C were introduced to eliminate *BsaI* and *BpiI* sites, respectively. The CDS were cloned into the Golden Gate level 0 vectors pAGM1287 and pAGM1299 ^84^. OPR fusions to YFP were generated by in-frame assembly of the level 0 with YFP at the C-terminus connected by a Gly-Ser linker. The final cloning cassettes containing the CaMV *35S* promoter and *tOcs* terminator in addition to the Oleosin-RFP plant selection marker were assembled in the pAGM55171 vector.

Plant stable transformation was carried-out by floral dip with *Agrobacterium tumefaciens* strain *GV3101* according to ^85^. To circumvent the *opr2opr3* mutant sterility, MeJA was applied to flower buds from 48 hours post-dipping until the first silique formation. Transformant selection relied on the seed coat RFP marker and single-insertion lines were identified by segregation analysis determining the ratio of fluorescent to non-fluorescent seeds.

### Subcellular targeting of the OPRs

Organelle variants of OPR3, OPR2, and OPR1 were generated by adding organelle targeting signals to the N- and/or C-terminal regions of the OPRs and/or YFP sequences (Supplementary Tab. 5). For nuclear targeting, OPR3ΔSRL was fused to the nuclear localization signal from Simian Virus 40 ^86,87^, while its targeting to the cytosol relied on a C-terminal nuclear export signal from the HIV1 REV ^88^. ER-localized OPR3ΔSRL was generated by flanking the sequence between an N-terminal signal peptide from the AtCAPE3 ^89^ and a C-terminal HDEL retention signal ^90^. Import to the chloroplast stroma and mitochondria, relied on an N-terminal target peptide from the AtRECA1 ^91^ and the Rieske protein ^92^, respectively.

### Transient expression in *N. benthamiana* leaves and protoplasts

*A. tumefaciens* strain *GV3101*, harboring OPR constructs, was cultured for 48 hours in LB medium with the appropriate antibiotics. The bacteria were syringe-infiltrated into *N. benthamiana* leaves according to ^93^. Leaf disc imaging was performed 48-72 hours post-infiltration.

Protoplasts were isolated from leaves of 4-week-old *N. benthamiana* plants, as described ^94^. Isolated protoplasts were co-transfected with level 1 plasmids of OPR3 constructs and organelle markers ^50^ using a mixture of 200 µl of protoplast suspension and 10 µg of DNA. Protoplasts were incubated at room temperature overnight after PEG-transformation and used for imaging.

### Confocal microscopy

Confocal microscopy utilized LSM880 and LSM900 laser scanning microscopes (Zeiss Germany, www.zeiss.de). Fluorophores were excited with 514 nm (YFP) and 561 nm (mCherry) laser lines and detected at 510–560 nm and 570-650 nm, respectively. Image acquisition and processing were performed using ZEN software (version 3.4, 2021, Zeiss).

### Phytohormone measurements

Measurements of OPDA, *dn*-OPDA, JA, and JA-Ile were performed using a standardized Ultra-performance liquid chromatography–tandem Mass Spectrometry (UPLC– MS/MS)-based method ^95^. 50 mg of powdered frozen tissue were extracted with 500 µl 100% LC-MS methanol supplemented with 50 ng of [^2^H_5_]OPDA, [^2^H_6_]JA, and [^2^H_2_]JA-Ile each as internal standards. After solid-phase extraction on HR-XC (Chromabond, Macherey-Nagel, www.mn-net.com), 10 µl of the eluate were analyzed via UPLC–MS/MS, and analyte content was determined relative to the internal standard peak heights. *dn*-OPDA content was determined using [^2^H_5_]OPDA as the internal standard.

### Determination of enzymatic activity of OPR3

OPR3 CDS inserted in pET28a(+) with *EcoR*I and *Not*I cloning sites was ordered from BioCat GmbH (www.biocat.com) and transformed into *Escherichia coli* BL21 Star (DE3) cells (ThermoFisher Scientific) by heat shock. Protein expression was performed in auto-induction medium at 16°C for three days ^96^. Harvested cells were resolved in buffer HisA (50 mM Tris/HCl pH 7.8, 100 mM NaCl, 10% (v/v) glycerol, 2 mM MgCl_2_ and 5 mM imidazole) supplemented with lysozyme, DNAse I and 0.2 mM PMSF. After a 30-minute incubation on ice, cells were sonicated and cleared lysate was applied to 1 mL HisTrap FastFlow column (Cytiva, www.cytivalifesciences.com). Protein was eluted with buffer HisB (50 mM Tris/HCl pH 7.8, 100 mM NaCl, 10% (v/v) glycerol, 2 mM MgCl_2_ and 500 mM imidazole), which was changed to storage buffer (50 mM Tris/HCl pH 7.8, 100 mM NaCl, 10% (v/v) glycerol, 2 mM MgCl_2_) by dialysis (MWCO 12-14 kDa) at 4°C overnight. Following SDS-PAGE, protein fractions were used for *in vitro* assays at 0.1 mg/mL together with 0.05 mM substrate, 1 mM NADPH and 10 mM NaCl in 20 mM Tris/HCl pH 7.8. Reactions were incubated at 30°C for 10 minutes and stopped by one volume acetonitrile. Enzyme deactivated by one volume acetonitrile was used as control. Assays were analyzed via UHPLC-high resolution (HR)-MS according to ^97^. Data was analyzed using the MassHunter Qualitative Analysis 10.0 software (Agilent Technologies).

## Figures and statistical analysis

Micrographs and photographs were processed through PHOTOSHOP 12.0.4 (Adobe Systems, http://www.adobe.com). The statistical analyses applied to the different datasets were performed using GraphPad Prism (www.graphpad.com).

## Data availability

The datasets generated and/or analyzed during this study are available from the corresponding author on request. Transcriptomic raw data were deposited in Sequence Read Archive (SRA) of NCBI (PRJNA1088739).

## Author contributions

Conceptualization, B.H and K.M.; Methodology, K.M, R.B, and F.S.; Investigation, K.M., R.B., F.S., and M.K.; Analyzing Data, K.M., I.F. and B.H.; Writing – Original Draft, K.M.; Writing – Review & Editing, B.H., K.M., and I.F.; Funding Acquisition, B.H. and I.F.;

## Acknowledgments

We thank Hagen Stellmach (IPB Halle, Germany) for help in jasmonate quantification, Andreas Schaller (University of Hohenheim, Germany) and Mats Hamberg (Karolinska Institute, Sweden) for providing seeds of *opr2opr3* and 4,5-ddh-JA, respectively. We thank Sylvestre Marillonnet (IPB Halle) for providing the Golden Gate modules and vectors used in this study. Khabat Vahabi (IPB Halle) is acknowledged for assistance in RNAseq preparation and data analysis. Claus Wasternack (IPB Halle) and Alain Tissier (IPB Halle) are acknowledged for critical reading of the manuscript. We thank Prof. Minoru Ueda (Tokhoku University, Japan) for sharing his data and helpful discussions on our manuscript.

## Supplementary information

**Fig. S1**: Seedlings of *opr2opr3* mutant show diminished AOC proteins content and produce lower OPDA levels compared to wild-type (Col-0).

**Fig. S2**: The experimental setup for transcriptome analyses.

**Fig. S3**: Hierarchical clustering and principal component analysis (PCA) illustrating the wound-induced transcriptional changes in Col-0, *opr2opr3* and *aos*.

**Fig. S4**: Wound-induced gene expression and hormone levels in *coi1* mutant leaves.

**Fig. S5:** The JA-dependent wound response occurring in wild-type seedlings.

**Fig. S6**: Transcriptome comparisons of wounded rosettes show limited signaling of OPDA.

**Fig. S7**: Application of OPDA results in the induction of JA-Ile signaling.

**Fig. S8**: The concept of the trans-organellar complementation of the *opr2opr3* mutant.

**Fig. S9**: OPR3 targeted to the ER, cytosol, nucleus, and chloroplasts colocalizes with the corresponding organelle markers in *N. benthamiana* protoplasts.

**Fig. S10**: Expression levels of the transgene in *opr2op3* mutants transformed with diverse OPR constructs.

**Fig. S11**: OPR3 reduces OPDA and 4,5-ddh-JA *in vitro*.

**Fig. S12:** OPDA levels at 1 h after wounding of seedlings of wild-type (WT) or *opr2opr3* transformed with either empty vector (EV), *35S::OPR2* or *35S::OPR1* targeted to peroxisomes (Px), cytosol (Cyt) or mitochondria (Mt).

**Tab. S1:** OPDA and JA levels shown in Fig. S1c-d.

**Tab. S2:** OPDA and JA levels shown in Fig. 1e.

**Tab. S3**: OPDA and JA levels shown in Fig. S5a.

**Tab. S4:** Genes selected from Taki et al. (2005) showing a similar wound-induced induction or repression in Col-0, *opr2opr3* and *aos*.

**Tab. S5:** Peptide sequences of the targeting signals used for subcellular targeting.

**Tab. S6:** Primers sequences used for mutants genotyping and RT-qPCR.

**Tab. S7:** Primer sequences used for Golden Gate cloning of OPR3, OPR2 and OPR1 and their subcellular targeting.

**Additional Supplementary Files**: Datasets 1-3

