## supplementary information for "Transcriptomics and trans-organellar complementation reveal a limited signaling capacity of 12-*cis*-oxo-phytodienoic acid in wounded Arabidopsis"

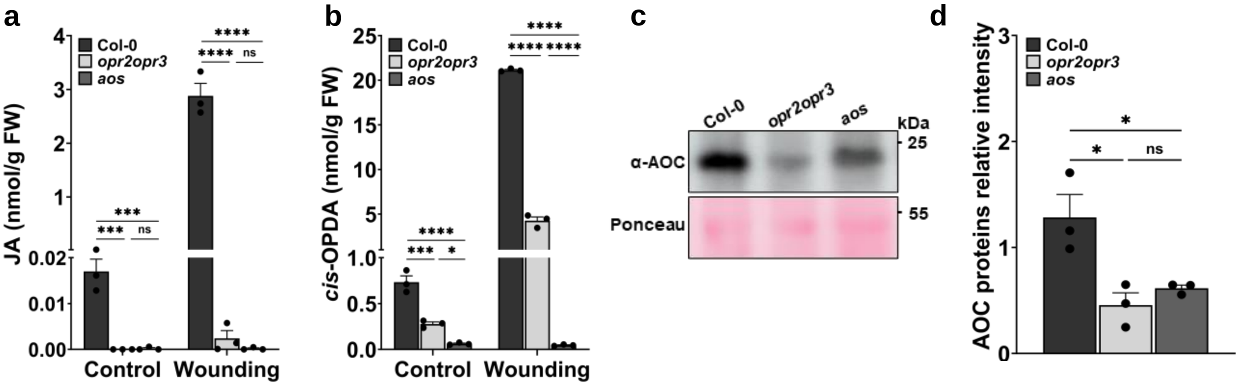


**Fig. S1: Seedlings of *opr2opr3* mutant show diminished AOC proteins content and produce lower *cis*-OPDA levels compared to wild-type (Col-0).**

**(a)** JA and **(b)** *cis*-OPDA levels in seedlings of wild type, *opr2opr3* and *aos* at control condition and at 1 h after wounding with forceps. For detailed values see Supplementary Tab. 1.

**(c)** AOC proteins content determination by immunoblot and **(d)** band intensity quantification relative to total protein loading stained by Ponceau S.

Bars in (a, b, d) represent means of three biological replicates with 30 seedlings each (single dots) ±SEM. Statistical significant differences among genotypes within each condition were calculated using One-Way ANOVA followed by Tukey-HSD and are indicated by asterisks with *p<0.05, **p<0.01, ***p<0.001, and ****p<0.0001.

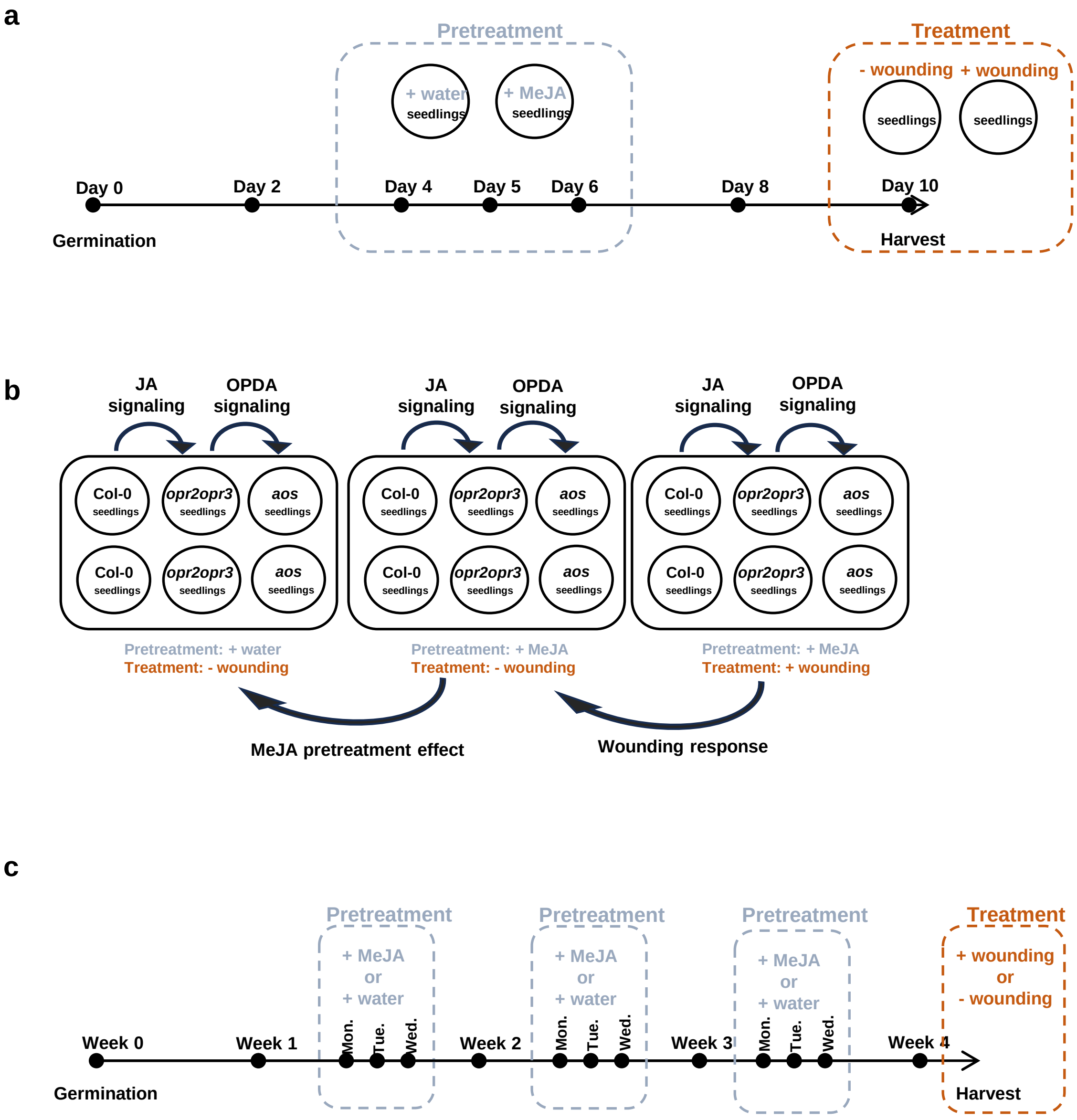


**Fig. S2: The experimental setup for transcriptome analyses.**
**(a)** Pretreatment and wounding of seedlings. Seedlings of wild type, *opr2opr3* and *aos* were grown in liquid MS medium. As pretreatment, 1 µM MeJA was applied at day 4, 5 and 6, and water was added as mock treatment. Wounding was performed using forceps at day 10. Wounded samples were harvested 1 hour later while the controls were kept unwounded. RNA was extracted and sent for mRNA sequencing to Novogene. Experiments were performed in triplicates with 120 seedlings per biological replicate.
**(b)** Different comparisons between each of the genotypes/conditions for transcriptome analyses. The comparisons of the transcriptomes between genotypes and/or conditions are shown in black arrows and indicate the signaling process to be characterized.
**(c)** Pretreatment and wounding of adult plants from wild type, *opr2opr3* and *aos* grown on soil. Pretreatment by spraying of the plants with 1 µM MeJA or water (as mock treatment) was performed every Monday, Tuesday and Wednesday starting within week two until week four of development. Wounding of the rosettes was performed after completion of the fourth week using forceps while controls were kept unwounded. Experiments were performed in triplicates with three rosettes pooled per biological replicate.

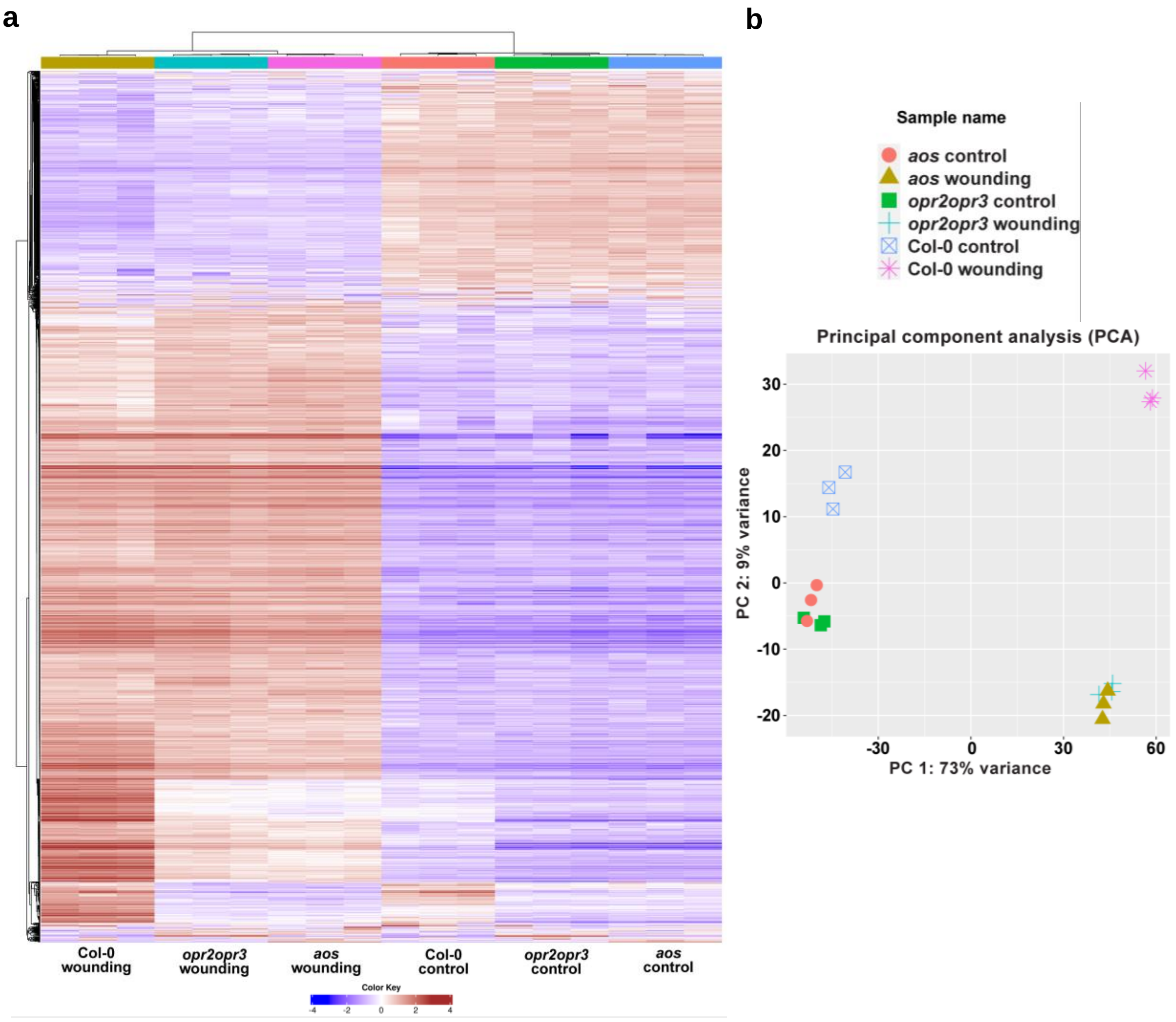


**Fig. S3: Hierarchical clustering and principal component analysis (PCA) illustrating the wound-induced transcriptional changes in Col-0, *opr2opr3* and *aos*.**

Ten-day old seedlings from Col-0, *opr2opr3* and *aos* were pretreated with MeJA during development and were either unwounded (control) or harvested 1 h after wounding with forceps (wounding).

**(a)** Heatmap with hierarchical clustering of the 2000 most variable genes based on FPKM values transformed into row-normalized Z-score with a threshold of 4. The data is centered by subtracting the average expression level for each gene. The distance matrix is 1- r, where r is Pearson's correlation coefficient. The average linkage is used (Bottom 25% of genes regarding expression level are excluded). Regions marked with blue brackets show a different expression pattern between the genotypes.

**(b)** PCA projection in two-dimensional space of the transcriptomic data with PC1 showing variation in the dataset due to wounding, while PC2 shows the variation related to the genotypes.

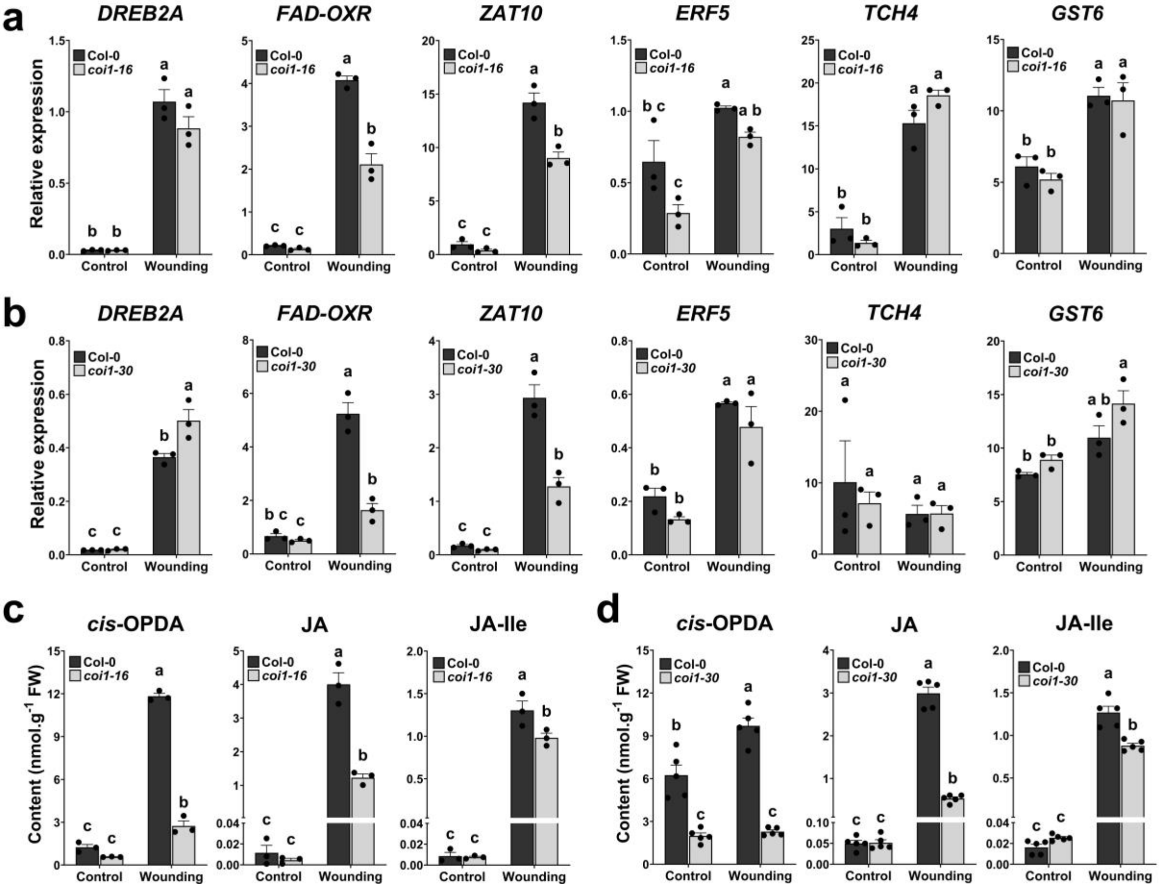


**Fig. S4: Wound-induced gene expression and hormone levels in *coi1* mutant leaves.**

Col-0, *coi1-16* and *coi1-30* plants were grown on soil. Leaves of 4-week-old plants remained either unwounded (Control) or were wounded with forceps and harvested 1 h post wounding (Wounding).

(**a**) RT-qPCR validation of the wound-induced up-regulation of *DREB2A2*, *FAD-OXR*, *ZAT10, ERF5, TCH4* and *GST6* in *coi1-16* leaves in comparison to wild type leaves (Col-0).

(**b**) RT-qPCR validation of the wound-induced up-regulation of *DREB2A2*, *FAD-OXR*, *ZAT10, ERF5, TCH4* and *GST6* in *coi1-30* leaves in comparison to wild type leaves (Col-0).

**(c-d)** Levels of *cis*-OPDA, *dn*-OPDA, JA and JA-Ile in adult rosettes of wild-type (Col-0) and *coi1-16* (c) or Col-0 and *coi1-30* (d) at control condition and 1 h after wounding.

Bars represent means of three biological replicates in (a-c) and five biological replicates in (d) (single dots; ±SEM). Statistically significant differences among treatments and genotypes were calculated using Two-Way ANOVA followed by Tukey-HSD and are indicated by different letters.

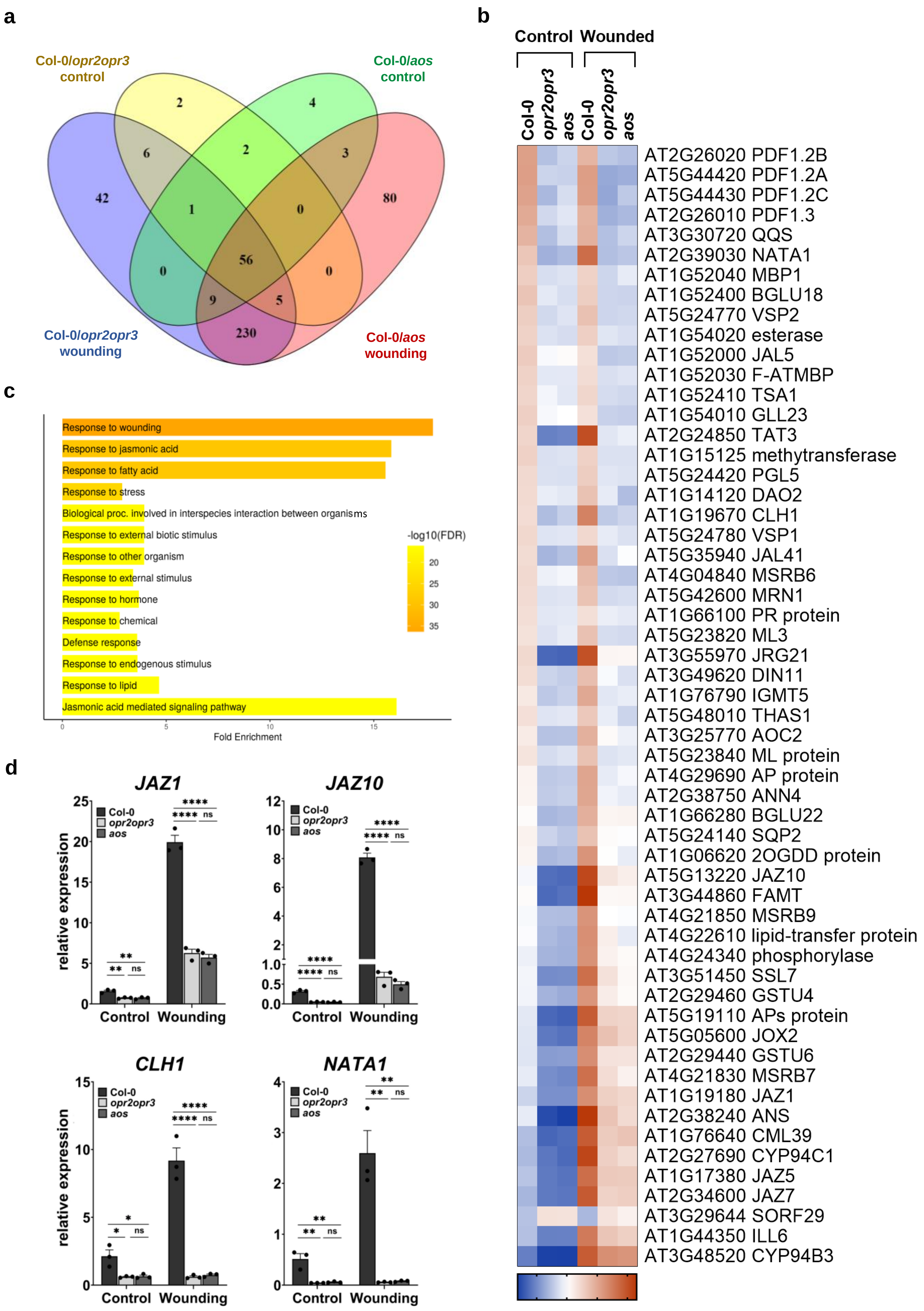
a
Col-0/opr2opr3 control
Col-0/aos control
Col-0/opr2opr3 wounding
Col-0/aos wounding
2
4
6
2
3
42
1
0
80
56
0
0
9
5
230
b
Control Wounded
Col-0
opr2opr3
aos
Col-0
opr2opr3
aos
AT2G26020 PDF1.2B
AT5G44420 PDF1.2A
AT5G44430 PDF1.2C
AT2G26010 PDF1.3
AT3G30720 QQS
AT2G39030 NATA1
AT1G52040 MBP1
AT1G52400 BGLU18
AT5G24770 VSP2
AT1G54020 esterase
AT1G52000 JAL5
AT1G52030 F-ATMBP
AT1G52410 TSA1
AT1G54010 GLL23
AT2G24850 TAT3
AT1G15125 methytransferase
AT5G24420 PGL5
AT1G14120 DAO2
AT1G19670 CLH1
AT5G24780 VSP1
AT5G35940 JAL41
AT4G04840 MSRB6
AT5G42600 MRN1
AT1G66100 PR protein
AT5G23820 ML3
AT3G55970 JRG21
AT3G49620 DIN11
AT1G76790 IGMT5
AT5G48010 THAS1
AT3G25770 AOC2
AT5G23840 ML protein
AT4G29690 AP protein
AT2G38750 ANN4
AT1G66280 BGLU22
AT5G24140 SQP2
AT1G06620 2OGDD protein
AT5G13220 JAZ10
AT3G44860 FAMT
AT4G21850 MSRB9
AT4G22610 lipid-transfer protein
AT4G24340 phosphorylase
AT3G51450 SSL7
AT2G29460 GSTU4
AT5G19110 APs protein
AT5G05600 JOX2
AT2G29440 GSTU6
AT4G21830 MSRB7
AT1G19180 JAZ1
AT2G38240 ANS
AT1G76640 CML39
AT2G27690 CYP94C1
AT1G17380 JAZ5
AT2G34600 JAZ7
AT3G29644 SORF29
AT1G44350 ILL6
AT3G48520 CYP94B3
-2 0 2 4
c
Response to wounding
Response to jasmonic acid
Response to fatty acid
Response to stress
Biological proc. involved in interspecies interaction between organisms
Response to external biotic stimulus
Response to other organism
Response to external stimulus
Response to hormone
Response to chemical
Defense response
Response to endogenous stimulus
Response to lipid
Jasmonic acid mediated signaling pathway
-log10(FDR)
20
25
30
35
0
5
10
15
Fold Enrichment
d
JAZ1
JAZ10
CLH1
NATA1
Col-0
opr2opr3
aos
relative expression
Control
Wounding
****
**
ns
*

**Fig. S5: The JA-dependent wound response occurring in wild type seedlings.**
Ten-day old seedlings from Col-0, *opr2opr3* and *aos* were pretreated with MeJA during development and were either unwounded (Control) or harvested 1 h after wounding with forceps (Wounding).
**(a)** Venn diagram representing differentially expressed genes (DEGs) identified by RNAseq in Col-0 compared to either *opr2opr3* or *aos* under control and wounding conditions (FDR cutoff = 0.05 and FC cutoff = 2).
**(b)** Heatmap illustrating the 56 JA-dependent DEGs common to all comparisons in **(a)**. Average FPKM values of three independent biological replicates were transformed into row normalized Z-Score.
**(c)** Bar plot showing Gene Ontology enrichment analysis summarizing the significant biological processes (FDR=0.05) enriched in Col-0 compared to the *opr2opr3* and *aos* mutants, with bars indicating gene fold enrichment and color scale indicating FDR values.
**(d)** RT-qPCR validation of selected JA-dependent DEGs from **(b)**. Transcript accumulation of *JAZ1*, *JAZ10*, *CLH1* and *NATA1* in Col-0, *opr2opr3* and *aos* at control and wounding conditions. Transcript levels were normalized to those of *PP2A3* (single dots; ±SEM). Statistically significant differences among genotypes within each condition were calculated using Two-Way ANOVA followed by Tukey-HSD and are indicated by asterisks with *$p<0.05$, **$p<0.01$, ***$p<0.001$, and ****$p<0.0001$.

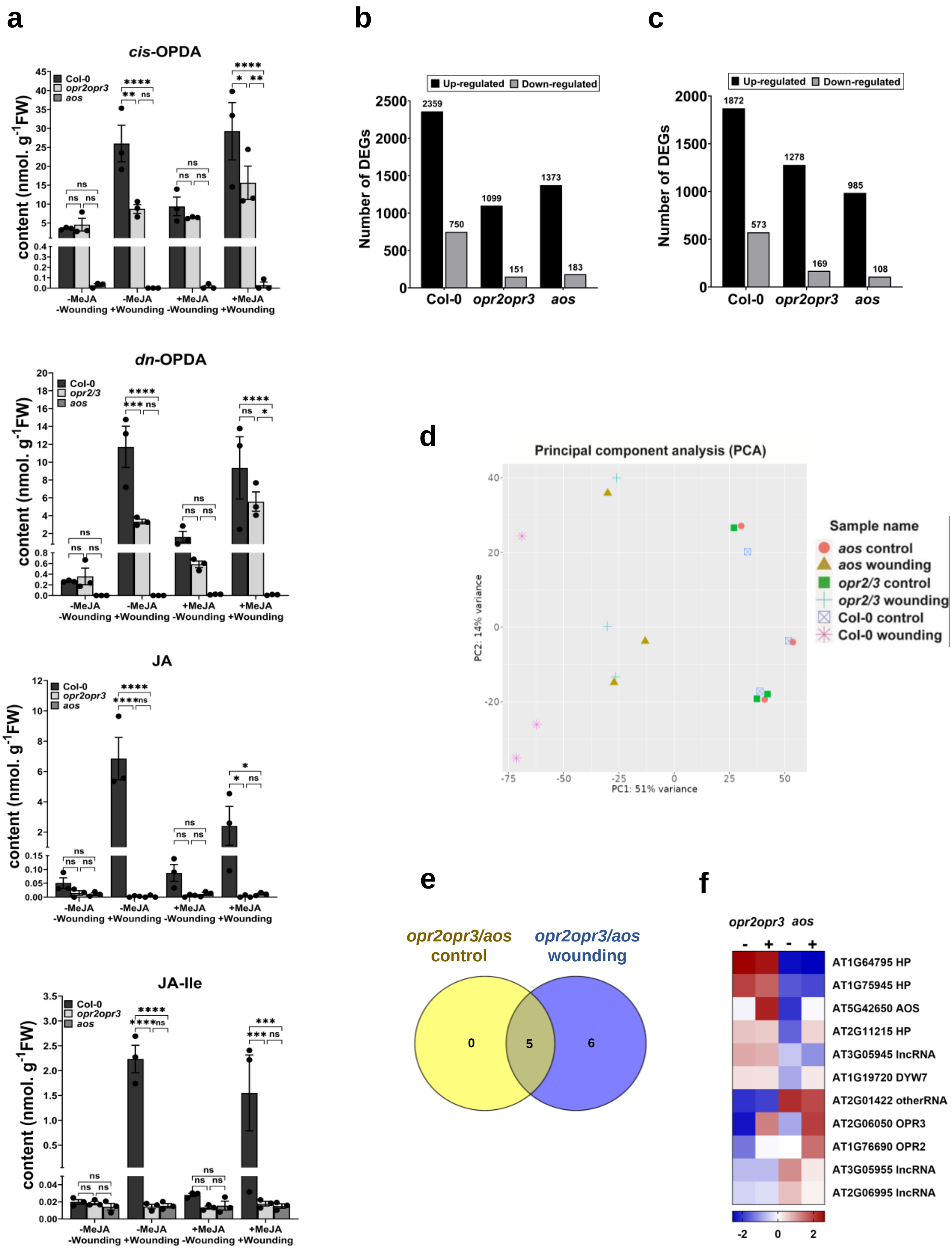
a
b
c
cis-OPDA
Col-0
opr2opr3
aos
content (nmol. g-1FW)
-MeJA -Wounding
-MeJA +Wounding
+MeJA -Wounding
+MeJA +Wounding
dn-OPDA
opr2/3
JA
JA-Ile
Up-regulated
Down-regulated
Number of DEGs
2359
750
1099
151
1373
183
1872
573
1278
169
985
108
d
Principal component analysis (PCA)
PC1: 51% variance
PC2: 14% variance
Sample name
aos control
aos wounding
opr2/3 control
opr2/3 wounding
Col-0 control
Col-0 wounding
e
opr2opr3/aos control
opr2opr3/aos wounding
0
5
6
f
opr2opr3
aos
AT1G64795 HP
AT1G75945 HP
AT5G42650 AOS
AT2G11215 HP
AT3G05945 lncRNA
AT1G19720 DYW7
AT2G01422 otherRNA
AT2G06050 OPR3
AT1G76690 OPR2
AT3G05955 lncRNA
AT2G06995 lncRNA

**Fig. S6: Transcriptome comparisons of wounded rosettes show limited signaling of OPDA.** Col-0, *opr2opr3* and *aos* plants grown on soil were pre-treated with 1 mM MeJA (+MeJA) or water (-MeJA) during development (see Material and Methods and Supplementary Fig. S2 for details). Leaves of 4-week-old plants were either unwounded (control) or harvested 1 h post wounding with forceps.

**(a)** Levels of *cis*-OPDA, *dn*-OPDA, JA and JA-Ile in adult rosettes of wild-type (Col-0), *opr2opr3*, and *aos* at control condition (-wounding) and 1 h after wounding with forceps (+wounding). For detailed values see Supplementary Tab. 3. Statistical significant differences among genotypes within each condition were calculated using One-Way ANOVA followed by Tukey-HSD and are indicated by asterisks with *$p<0.05$, **$p<0.01$, ***$p<0.001$, and ****$p<0.0001$.

**(b)** Number of differentially regulated genes (DEGs) after wounding compared to non-wounded controls in seedlings of Col-0, *opr2opr3* and *aos*, all pretreated with water during development. DEGs were identified using FDR and FC cutoffs of 0.05 and 2, respectively.

**(c)** Number of DEGs after wounding compared to non-wounded controls in seedlings of Col-0, *opr2opr3* and *aos* mutants, all pretreated with 1 µM MeJA during development. DEGs were identified using FDR and FC cutoffs of 0.05 and 2, respectively.

**(d)** PCA projection in two-dimensional space of the transcriptome data at wounding and control conditions, with PC1 showing variation in the dataset due to wounding while PC2 showing the variation due to the genotypes. Venn diagram showing the wound response in Col-0 compared to *opr2opr3* and *aos* mutants which were pretreated with water during development. Differentially expressed genes (DEGs) after wounding compared to non-wounded control for each genotype showed a common response with 1125 DEGs. DEGs were identified using FDR and FC cutoffs of 0.05 and 2, respectively.

**(e)** Venn diagram showing a total of 11 DEGs in *opr2opr3* compared to *aos* at control and wounding conditions, with 5 DEGs being common to both conditions. (FDR cutoff = 0.05 and FC cutoff = 2).

**(f)** Heatmap illustrating the DEGs from (C). Average FPKM values of three independent biological replicates were transformed into normalized Z-Score.

**a**

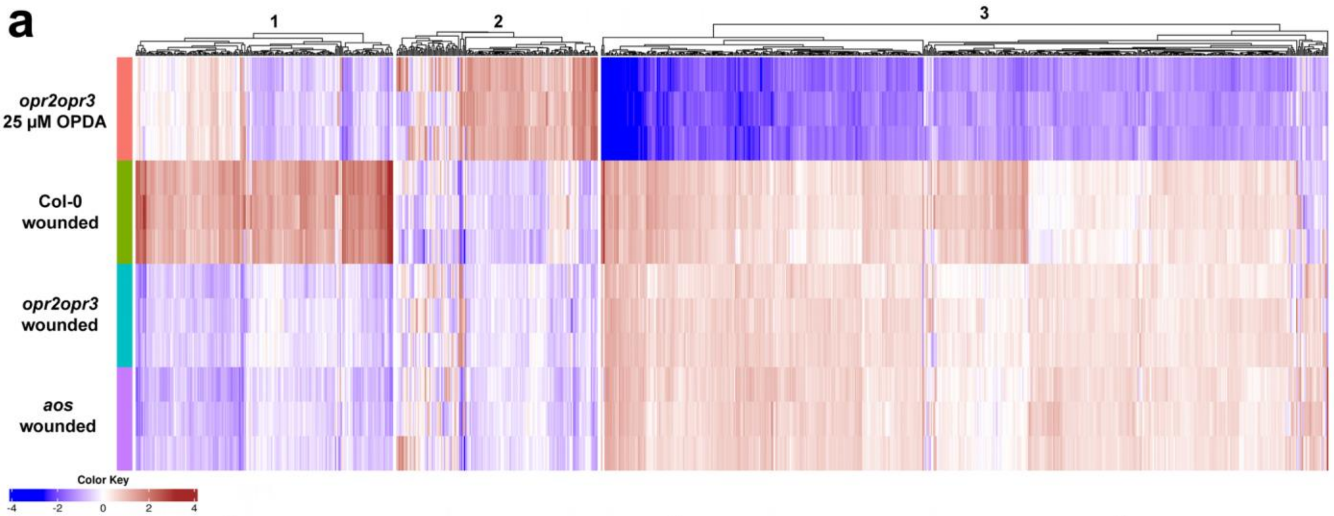


**b**

1

| FDR | nGenes | P.S | P.E | Pathway |
|---|---|---|---|---|
| 8.31E-28 | 33 | 222 | 16.14 | Response to wounding |
| 1.44E-25 | 30 | 196 | 16.42 | Response to jasmonic acid |
| 1.81E-25 | 30 | 200 | 16.08 | Response to fatty acid |
| 3.09E-15 | 83 | 3906 | 2.41 | Response to stress |
| 5.10E-15 | 17 | 98 | 18.32 | Jasmonic acid mediated signaling pathway |
| 5.10E-15 | 13 | 41 | 33.49 | Regulation of jasmonic acid mediated signaling pathway |
| 9.82E-15 | 17 | 103 | 17.43 | Cellular response to jasmonic acid stimulus |
| 1.68E-14 | 17 | 107 | 16.78 | Cellular response to fatty acid |
| 1.42E-11 | 46 | 1700 | 3.27 | Defense response |
| 1.42E-11 | 42 | 1457 | 3.56 | Response to external biotic stimulus |

2

| FDR | nGenes | P.S | P.E | Pathway |
|---|---|---|---|---|
| 8.07E-05 | 3 | 3 | 203.29 | Sulfate reduction |
| 1.28E-04 | 28 | 2383 | 2.6 | Regulation of transcription, DNA-templated |
| 1.28E-04 | 33 | 3140 | 2.31 | Regulation of nitrogen compound metabolic process |
| 1.28E-04 | 28 | 2383 | 2.6 | Regulation of nucleic acid-templated transcription |
| 1.28E-04 | 28 | 2383 | 2.6 | Regulation of RNA biosynthetic process |
| 1.57E-04 | 29 | 2587 | 2.47 | Regulation of nucleobase-containing compound metabolic process |
| 1.57E-04 | 33 | 3210 | 2.26 | Regulation of primary metabolic process |
| 2.14E-04 | 4 | 22 | 38.72 | Sulfate assimilation |
| 2.14E-04 | 9 | 260 | 7.59 | Phosphorelay signal transduction system |
| 2.14E-04 | 28 | 2555 | 2.43 | Transcription, DNA-templated |

3

| FDR | nGenes | P.S | P.E | Pathway |
|---|---|---|---|---|
| 6.37E-82 | 91 | 241 | 15.25 | Cellular response to decreased oxygen levels |
| 6.37E-82 | 91 | 241 | 15.25 | Cellular response to oxygen levels |
| 6.37E-82 | 91 | 239 | 15.38 | Cellular response to hypoxia |
| 1.93E-80 | 93 | 265 | 14.17 | Response to hypoxia |
| 7.93E-80 | 93 | 269 | 13.96 | Response to decreased oxygen levels |
| 9.91E-80 | 93 | 270 | 13.9 | Response to oxygen levels |
| 5.97E-60 | 245 | 3906 | 2.71 | Response to stress |
| 1.01E-42 | 167 | 2214 | 3.09 | Response to abiotic stimulus |
| 1.37E-42 | 200 | 3152 | 2.63 | Response to chemical |
| 8.10E-36 | 116 | 1265 | 3.8 | Cellular response to stress |

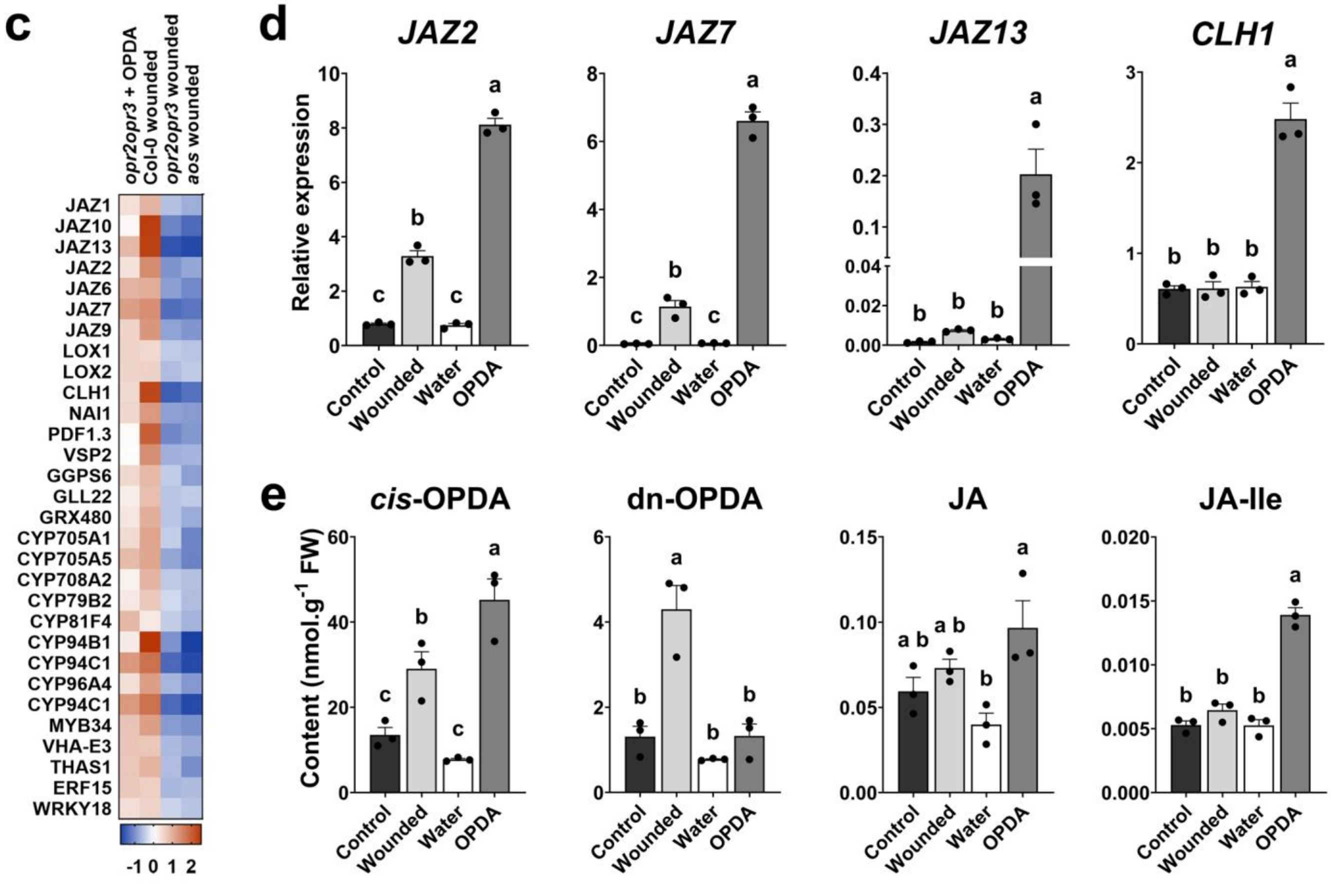

**Fig. S7: Application of OPDA results in the induction of JA-Ile signaling.**

**(a)** Heat map of DEGs comparing the transcriptional response of *opr2opr3* seedlings upon treatment with OPDA with the response of seedlings from Col-0, *opr2opr3* and *aos* upon wounding. K-means clustering of the 1000 most variable genes across the RNA-seq samples using FPKM values transformed into row-normalized Z-score of three biological replicates.

**(b)** Gene ontology (GO) enrichment analysis of each subcluster using False discovery rate (FDR), pathway size (P.S), and fold enrichment (F.E) as enrichment parameters. GO was performed based on biological processes.

**(c)** Heatmap of DEGs showing enrichment in JA signaling pathway genes and genes involved in sulfur assimilation in seedlings of *opr2opr3* following application of 25 µM OPDA for 30 min. DEGs were plotted and compared to those from wounded seedlings of Col-0, *opr2opr3* and *aos* using average FPKM values transformed into row-normalized Z-score of three biological replicates.

**(d)** RT-qPCR validation of the up-regulation of JA-responsive genes *JAZ2*, *JAZ7*, *JAZ13* and *CLH1* in *opr2opr3* seedlings treated with OPDA in comparison to wounded *opr2opr3* seedlings. OPDA-treated and wounded *opr2opr3* seedlings were compared to their controls, non-wounded (control) and water-treated seedlings, respectively.

**(e)** *cis*-OPDA, dn-OPDA, JA and JA-Ile accumulation in *opr2opr3* seedlings treated with 25 µM OPDA in comparison to wounded *opr2opr3* seedlings producing OPDA endogenously.

Bars (d-e) represent means of three biological replicates with 120 seedlings each (single dots; ±SEM). Statistically significant differences among treatments were calculated using Two-Way ANOVA followed by Tukey-HSD and are indicated by different letters.

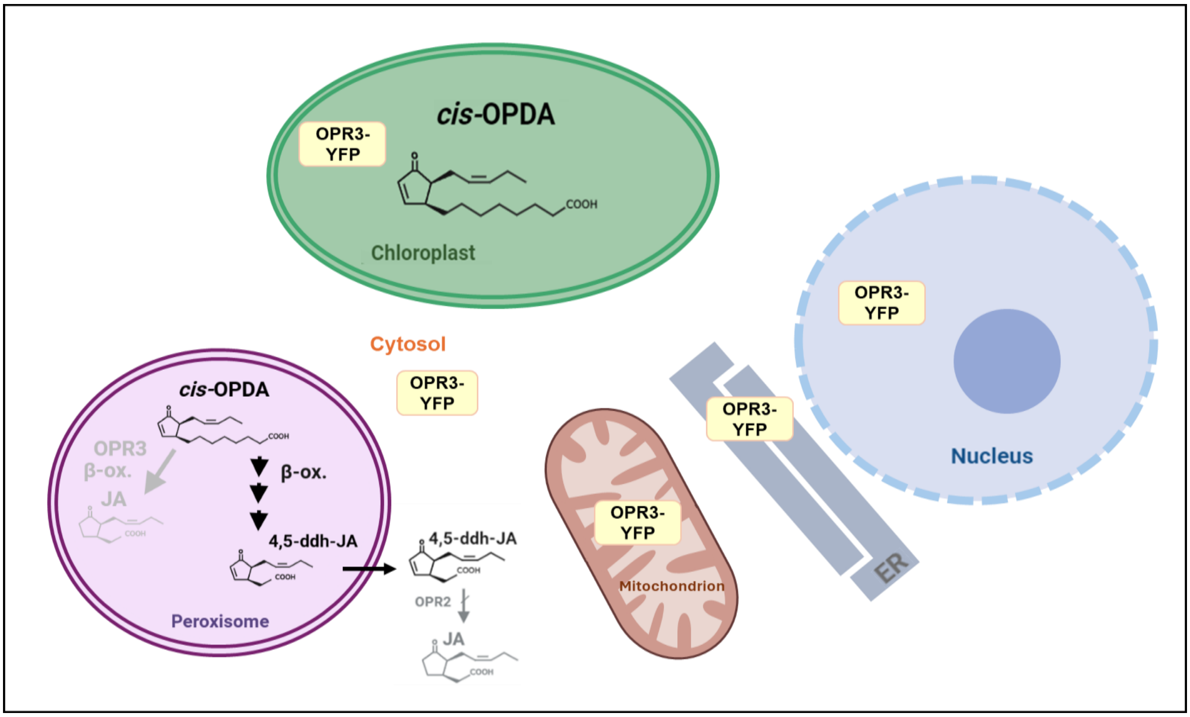


**Fig. S8: The concept of the trans-organellar complementation of the *opr2opr3* mutant.**
The *opr2opr3* mutant is complemented with OPR3 fused to YFP and targeted to a different organelle than peroxisomes.
Pathways in grey are those blocked in the *opr2opr3* mutant. The loss of function of OPR3 and OPR2 stops the formation of OPC-8/OPC-6 from *cis*-OPDA/*dn*-OPDA and JA from 4,5-ddh-JA, respectively. The question to be answered is, whether OPDA translocate to other organelles, where it will be converted by OPR3 to OPC-8 resulting in production of JA/JA-Ile.
Abbreviations: OPR3, 12-oxophytodienoate-10,11-reductase 3; OPR2, 12-oxophytodienoate-10,11-reductase 2; *cis*-(+)-OPDA, *cis*-(+)-12-oxophytodienoic acid; *dn*-OPDA, dinor-12-oxophytodienoic acid; 4,5-ddh-JA, 4,5-didehydrojasmonic acid; OPC-8, 3-oxo-2-(2-pentenyl)-cyclopentane-1-octanoic acid; OPC-6, 3-oxo-2-(2-pentenyl)-cyclopentane-1-hexanoic acid; JA, jasmonic acid.

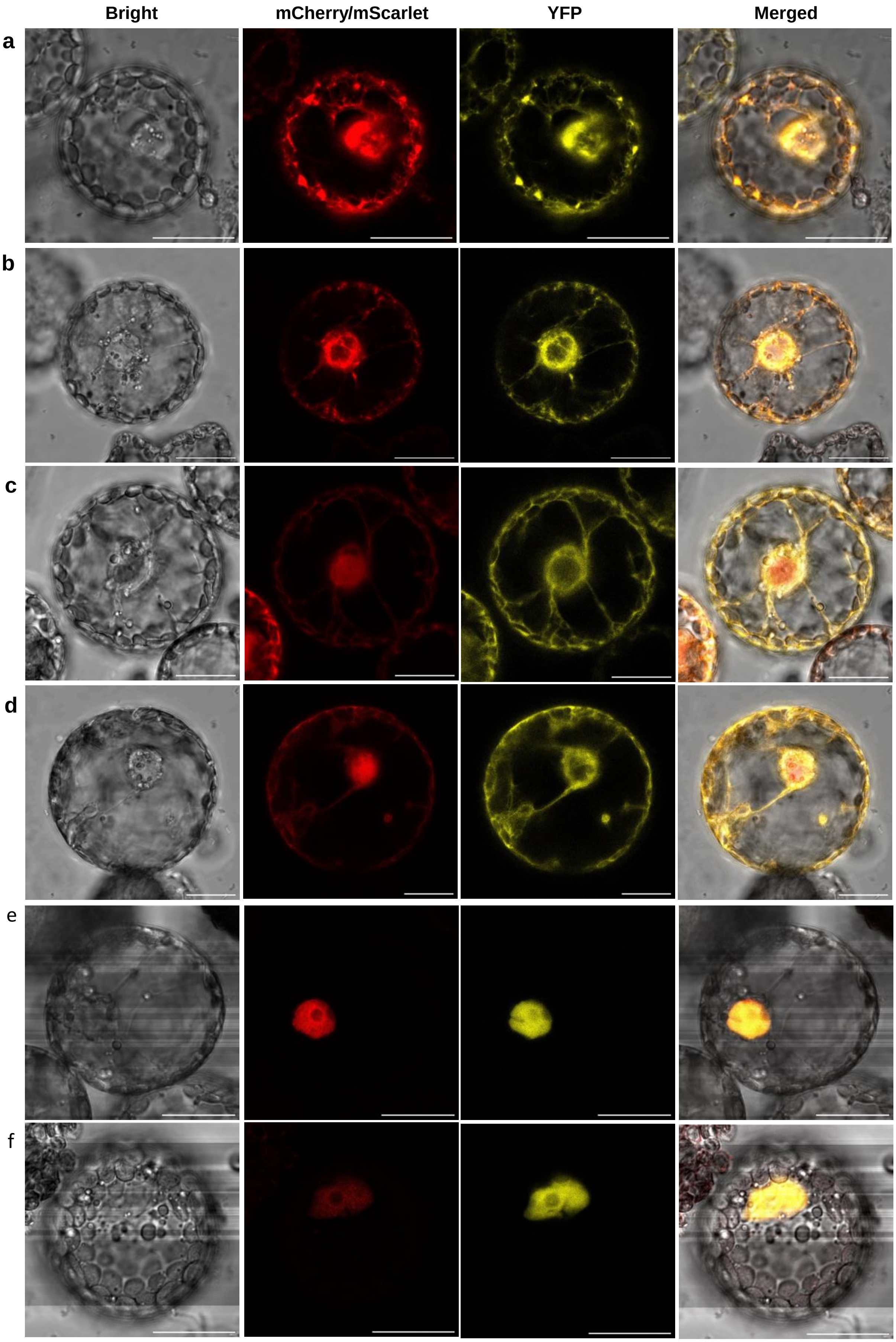
Bright
mCherry/mScarlet
YFP
Merged
a
b
c
d
e
f

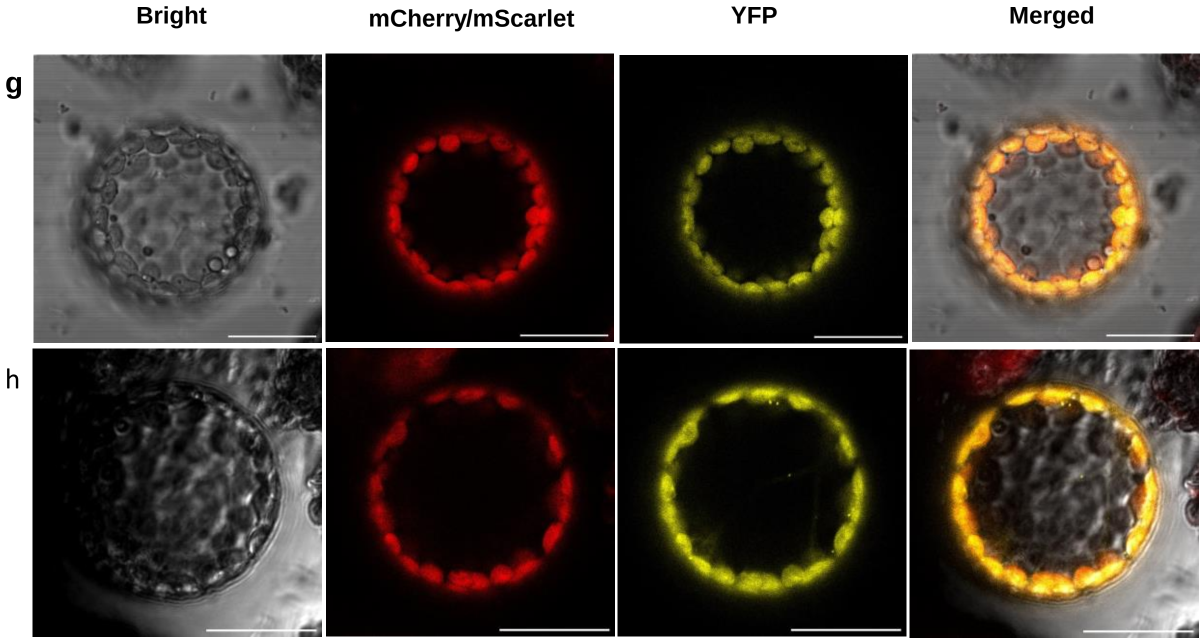


**Fig. S9: OPR3 targeted to the ER, cytosol, nucleus, and chloroplasts colocalizes with the corresponding organelle markers in *N. benthamiana* protoplasts.** The CaMV *35S* promoter and *tOCS* terminator were used to drive the expression of *OPR3ΔSRL-YFP* and *YFP-OPR3ΔSRL* for each organelle targeting. The protoplasts were transiently co-transformed with each construct and its corresponding organelle marker and imaged 12 h post transformation. The yellow signal corresponds to YFP, the red to the mCherry or mScarlet organelle marker, and the gray to the bright field. Scale bars: 20 µm.

**(a, b)** ER-targeted OPR3 colocalizes with the WAK-mCherry-KDEL marker. The YFP fusions being SIP-OPR3ΔSRL-YFP-HDEL (**a**) and SIP-YFP-OPR3ΔSRL-HDEL (**b**).

**(c, d)** Cytosol-targeted OPR3 colocalizes with the mScarlet marker in the cytosol but not inside the nucleus. The YFP fusions being OPR3ΔSRL-YFP-NES (**c**) and YFP-OPR3ΔSRL-NES (**d**).

**(e, f)** Nucleus-targeted OPR3 colocalizes with the HTA6-mCherry marker. The YFP fusions being NLS-OPR3ΔSRL-YFP (**e**) and NLS-YFP-OPR3ΔSRL (**f**).

**(g, h)** Plastid stroma-targeted OPR3 colocalizes with the RubisCO-mCherry marker. The YFP fusions being cTP-OPR3ΔSRL-YFP (**g**) and cTP-YFP-OPR3ΔSRL (h).

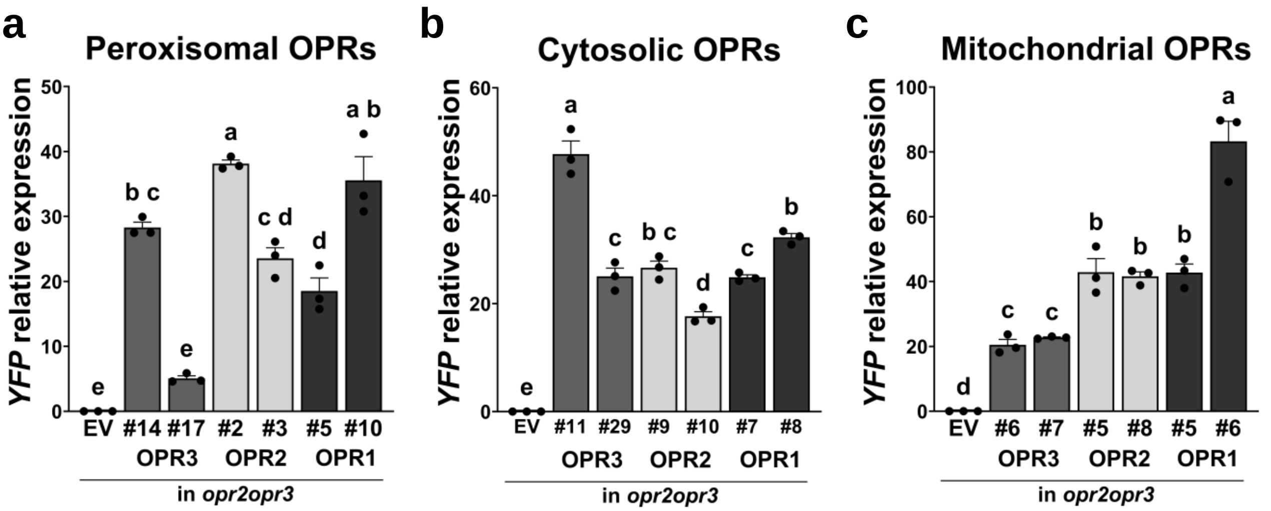


**Fig. S10: Expression levels of the transgene in *opr2opr3* mutants transformed with diverse OPR-YFP constructs.**

*YFP* expression levels in the *opr2opr3* complementation lines determined by RT-qPCR in relation to *PP2A3* (complementation constructs are listed in Supplementary table S5).

**(a)** *YFP* transgene expression levels in *opr2opr3* transformed with empty vector (EV) or the peroxisome-targeted OPR3-YFP, OPR2-YFP and OPR1-YFP.

**(b)** *YFP* transgene expression levels in the *opr2opr3* transformed with empty vector (EV) or the cytosol-targeted OPR3-YFP, OPR2-YFP and OPR1-YFP.

**(c)** *YFP* transgene expression levels in the *opr2opr3* transformed with empty vector (EV) or the mitochondria-targeted OPR3-YFP, OPR2-YFP and OPR1-YFP.

Numbers refer to independent transgenic lines, which were selected for single insertion and homozygosity. Bars represent means of three biological replicates with 30 seedlings each (single dots; ±SEM). Statistically significant differences were calculated using One-Way ANOVA followed by Tukey HSD ($p<0.05$) and are denoted by different letters.

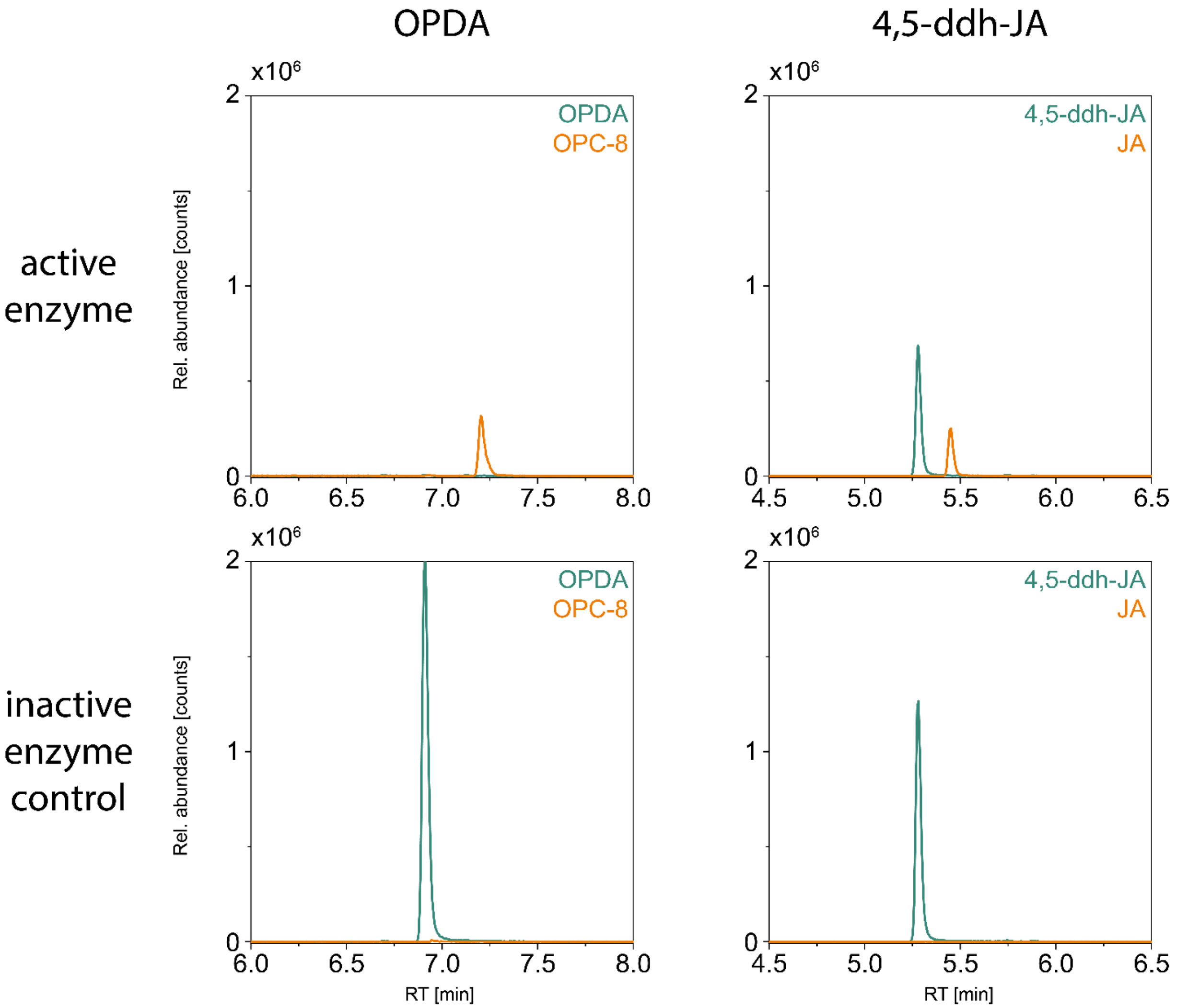


**Fig. S11: OPR3 reduces OPDA and 4,5-ddh-JA *in vitro*.**
Heterologous expressed and purified 6x-His-OPR3 was tested in *in vitro* assays at a concentration of 0.1 mg/ml. Either, 50µM OPDA or 4,5-ddh-JA were added to start the reaction and were incubated at 30 °C for 10 min while slowly shaking. Reactions were stopped by adding one volume of acetonitrile. Data collection was performed by UHPLC-HRMS in positive (OPDA) or negative (4,5-ddh-JA) electrospray ionization mode. Figure shows the extracted ion chromatograms (EICs) of OPDA ($[M+H]^+$ 293.211 Da) and OPC-8 ($[M+H]^+$ 295.227) or 4,5-ddh-JA ($[M-H]^-$ 207.103) and JA ($[M-H]^-$ 209.118) in the assays either with active or inactive enzyme. Identity of *in vitro* products was confirmed by authentic standards or MS/MS fragmentation. Figure is representative for four experiments.

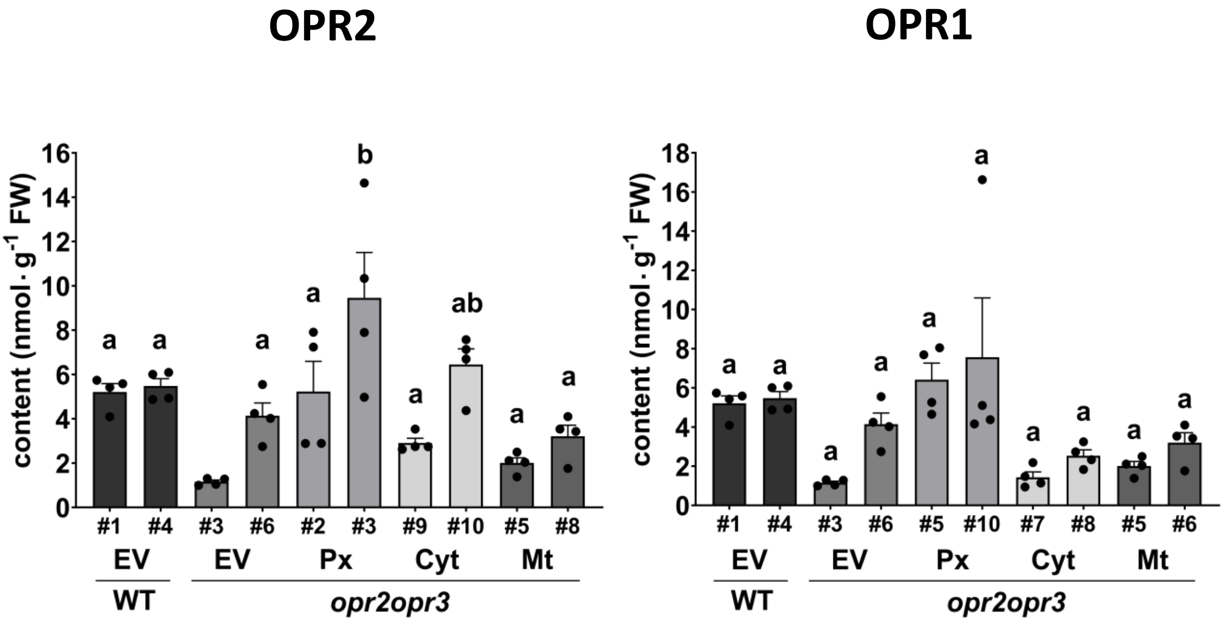


**Fig. S12: OPDA levels at 1 h after wounding of seedlings of wild type (WT) or *opr2opr3* transformed with either empty vector (EV), *35S::OPR2* or *35S::OPR1* targeted to peroxisomes (Px), cytosol (Cyt) or mitochondria (Mt).**
Numbers refer to independent transgenic lines, which were selected for single insertion and homozygosity. Bars represent means of four biological replicates with 120 seedlings each (single dots; ±SEM). Statistically significant differences between *opr2opr3* transformed with empty vector and transformed with the OPR3 constructs were calculated using One-Way ANOVA followed by Tukey HSD ($p<0.05$) and are denoted by different letters.

**Supplementary Tab. 1: OPDA and JA levels shown in Fig. S1c-d.**

| | ***cis*-OPDA** (nmol/g fresh weight) | | **JA** (nmol/g fresh weight) | |
|---|---|---|---|---|
| | Control | Wounding | Control | Wounding |
| Col-0 | 0.733 (± 0.119) | 21.127 (± 0.166) | 0.016 (± 0.0046) | 2.879 (± 0.400) |
| *opr2opr3* | 0.277 (± 0.038) | 4.251 (± 0.718) | 0 (± 0) | 0.00243 (± 0.0029) |
| *aos* | 0.060 (± 0.012) | 0.045 (± 0.008) | 0.00018 (± 0.00032) | 0.00015 (± 0.00027) |

**Supplementary Tab. 2: OPDA and JA levels shown in Fig. 1e.**

| | ***cis*-OPDA** (nmol/g fresh weight) | | | **JA** (nmol/g fresh weight) | | |
|---|---|---|---|---|---|---|
| | Mock -Wounding | MeJA -Wounding | MeJA +Wounding | Mock -Wounding | MeJA -Wounding | MeJA +Wounding |
| Col-0 | 9.585 (± 4.215) | 12.343 (± 2.897) | 26.857 (± 1.675) | 0.0484 (± 0.016) | 0.169 (± 0.051) | 5.253 (± 0.678) |
| *opr2opr3* | 2.2107 (± 0.540) | 13.522 (± 3.048) | 29.056 (± 6.894) | 0.00106 (± 0.00098) | 0.059 (± 0.0141) | 0.073 (± 0.009) |
| *aos* | 0.135 (± 0.1098) | 0.208 (±0.218) | 0.241 (±0.071) | 0.0013 (±0.0011) | 0.0536 (±0.0418) | 0.0655 (±0.0145) |

**Supplementary Tab. 3: OPDA and JA levels shown in Fig. S5a.**

| | ***cis*-OPDA** (nmol/g fresh weight) | | | |
|---|---|---|---|---|
| | - MeJA - Wounding | - MeJA + Wounding | + MeJA - Wounding | + MeJA + Wounding |
| Col-0 | 3.545 (± 0.3406) | 25.998 (±8.359) | 9.406892 (± 4.245) | 29.243 (± 13.116) |
| *opr2opr3* | 4.570 (± 2.895) | 8.731 (± 2.029) | 6.423 (± 0.2401) | 15.633 (± 7.648) |
| *aos* | 0.0277 (± 0.0242) | 0 (± 0) | 0.0140 (± 0.0244) | 0.0290 (± 0.0503) |
| | **JA** (nmol/g fresh weight) | | | |
| | - MeJA - Wounding | - MeJA + Wounding | + MeJA - Wounding | + MeJA + Wounding |
| Col-0 | 0.0506 (± 0.0331) | 6.849 (± 2.423) | 0.0873 (± 0.0527) | 2.406 (± 2.226) |
| *opr2opr3* | 0.0151 (± 0.0144) | 0.00286 (± 0.0025) | 0.00535 (± 0.0048) | 0.0026 (± 0.0046) |
| *aos* | 0.01088 (± 0.0079) | 0.0024 (± 0.0042) | 0.0118 (± 0.0090) | 0.0110 (± 0.00534) |

**Supplementary Tab. 4: Genes selected from Taki et al. 2005 showing a similar wound-induced induction or repression in Col-0, *opr2opr3* and *aos*.**

| AGI | Gene name | LogFC[A] | | | Adjusted p-value[B] | | | Description |
|---|---|---|---|---|---|---|---|---|
| | | Col-0 | *opr2-opr3* | *aos* | Col-0 | *opr2-opr3* | *aos* | |
| **Up regulated genes** | | | | | | | | |
| AT5G42380 | CML37 | 5.09 | 4.94 | 4.76 | 2.2E-13 | 5.3E-13 | 6.3E-13 | Calcium-binding protein CML37 |
| AT3G25250 | OXI1 | 3.76 | 3.84 | 3.53 | 1.9E-11 | 2.1E-11 | 4.9E-11 | Serine/threonine-protein kinase OXI1 |
| AT5G59820 | ZAT12 | 3.54 | 3.50 | 3.29 | 4.3E-10 | 6.3E-10 | 1.2E-09 | Zinc finger protein ZAT12 |
| AT5G35735 | HYP1 | 3.42 | 3.73 | 3.62 | 3.2E-11 | 1.7E-11 | 2.1E-11 | Cytochrome B561 |
| AT1G61340 | FBS1 | 3.40 | 3.44 | 3.52 | 1.2E-10 | 1.4E-10 | 1.0E-10 | F-BOX stress induced 1 |
| AT2G46400 | WRKY46 | 3.14 | 3.31 | 3.25 | 2.6E-09 | 1.9E-09 | 2.1E-09 | Probable WRKY transcription factor 46 |
| AT3G04640 | - | 3.11 | 3.01 | 3.11 | 9.7E-10 | 1.8E-09 | 1.2E-09 | Glycine-rich protein |
| AT2G22500 | PUMP5 | 2.89 | 2.74 | 2.82 | 1.4E-11 | 3.3E-11 | 2.2E-11 | Mitochondrial uncoupling protein 5 |
| AT3G46080 | ZAT8 | 2.53 | 2.55 | 2.34 | 1.3E-08 | 1.5E-08 | 3.6E-08 | Zinc finger protein ZAT8 |
| AT1G66090 | TN3 | 2.49 | 2.71 | 2.56 | 1.4E-08 | 6.4E-09 | 1.1E-08 | Disease resistance protein (TIR-NBS class) |
| AT4G24570 | DIC2 | 2.24 | 2.12 | 1.83 | 1.2E-09 | 2.9E-09 | 1.4E-08 | DICARBOXYLATE CARRIER 2 |
| AT5G27420 | ATL31 | 2.18 | 1.93 | 2.00 | 9.0E-08 | 4.0E-07 | 2.6E-07 | E3 ubiquitin-protein ligase ATL31 |
| AT4G24160 | - | 2.01 | 2.43 | 2.30 | 1.0E-10 | 1.8E-11 | 3.0E-11 | 1-acylglycerol-3-phosphate O-acyltransferase |
| AT5G57560 | XTH22 | 1.69 | 1.57 | 1.42 | 1.7E-08 | 4.6E-08 | 1.3E-07 | Xyloglucan endotransglucosylase/hydrolase |
| AT1G15520 | ABCG40 | 1.64 | 1.69 | 1.33 | 6.6E-07 | 5.4E-07 | 7.2E-06 | ABC transporter G family member 40 |
| AT5G54490 | PBP1 | 1.51 | 1.70 | 1.48 | 3.8E-07 | 1.2E-07 | 5.2E-07 | Pinoid-binding protein 1 |
| AT1G59660 | NUP98B | 1.49 | 1.20 | 1.44 | 2.0E-08 | 2.6E-07 | 3.2E-08 | Nuclear pore complex protein NUP98B |
| AT1G59860 | HSP17.6A | 1.48 | 1.25 | 1.20 | 6.4E-06 | 4.3E-05 | 6.0E-05 | 17.6 kDa class I heat shock protein 1 |
| AT1G16030 | HSP70-5 | 1.35 | 1.37 | 1.41 | 2.5E-04 | 2.6E-04 | 1.9E-04 | Heat shock protein 70-5 |
| AT5G45340 | CYP707A3 | 1.26 | 1.64 | 1.56 | 6.0E-04 | 6.2E-05 | 9.8E-05 | Abscisic acid 8'-hydroxylase 3 |
| **Down regulated genes** | | | | | | | | |
| AT1G78000 | SULTR1;2 | -1.70 | -1.35 | -1.01 | 7.4E-07 | 9.8E-06 | 1.7E-04 | Sulfate transporter 1.2 |

[A] $log_2$ fold change with a cutoff of 1 calculated in the respective genotypes at wounding in relation to the control.
[B] False discovery rate corrected p-value with a cutoff of 0.05

**Supplementary Tab. 5: Peptide sequences of the targeting signals used for subcellular targeting.**

| **Cell compartment** | **Targeting signal** | **Peptide sequence** | **Sequence position** | **Constructs used for *opr2opr3* complementation** |
|---|---|---|---|---|
| Peroxisome | Peroxisome targeting signal 1 (PTS1) | SRL (OPR3), SKL (OPR2-OPR1) | C-terminus | *35S::OPR3ΔSRL-YFP-SRL::tOcs*<br>*35S::OPR2-YFP-SKL::tOcs*<br>*35S::OPR1-YFP-SKL::tOcs* |
| Nucleus | Nuclear targeting signal (NLS) | MASSPPKKKRKVSWK | N-terminus | *35S::NLS-OPR3ΔSRL-YFP-NES::tOcs* |
| Cytosol | Nuclear exclusion signal (NES) | LQLPPLERLTLD (OPR3) | C-terminus | *35S::OPR3ΔSRL-YFP-NES::tOcs*<br>*35S::OPR2-YFP::tOcs*<br>*35S::OPR1-YFP::tOcs* |
| ER | Signal peptide (SP) | MASKSVVVFLFLALVASSVV | N-terminus | *35S::SP-OPR3ΔSRL-YFP-RS::tOcs* |
| ER | Retention signal (RS) | HDEL | C-terminus | |
| Chloroplast | Chloroplast targeting peptide (cTP) | MDSQLVLSLKLNPSFTPLSPLFPFTPC<br>SSFSPSLRFSSCYSRRLYSPVTVY | N-terminus | *35S::cTP-OPR3ΔSRL-YFP::tOcs* |
| Mitochondria | Mitochondria targeting peptide (mTP) | MMLRVAGRRLSSSAARSSSSFFTRSS<br>FTVTDDSSPARSPSPSLTSSFLDQIRG<br>FSSNSVSPAHQLGLVSDLPATVAAIKN<br>PSSKIVYDDSNHERYPPGDP | N-terminus | *35S::mTP-OPR3ΔSRL-YFP::tOcs*<br>*35S:: mTP-OPR2-YFP::tOcs*<br>*35S:: mTP-OPR1-YFP::tOcs* |

**Supplementary Tab. 6: Primers sequences used for mutants genotyping and RT-qPCR.**

| Genotyping primers | |
|---|---|
| Genotype | Primer sequence (5´->3´) |
| *aos (dde2-2)* | for_CGAACATGTAGAGCAGCAACAG |
| | rev_GCCAGAGTCTCCAATAGATCTC |
| *opr2opr3 (SK24765)* | for_AATCCGTGTAGCCAACAACTG |
| | rev_CAGCCACATTCAAAGAAAAGG |
| *aos (SALK_116381)* | for_GTGGGTTATTGCTGATCATCC |
| | rev_AGCTGTTGATTCAAGGGAAGG |
| *coi1-30* | LBb1.3 ATTTTGCCGATTTCGGAAC |
| | RP CTGCAGTGTGTAACGATGCTC |
| RT-qPCR primers | |
| Gene | Sequence 5´->3´ |
| *PP2A3* (AT1G13320) | for_AGACAAGGTTCACTCAATCCGTG |
| | rev_CATTCAGGACCAAACTCTTCAGC |
| *LOX2* (AT3G45140) | for_ACGCTCGTGCACGCCAAAGT |
| | rev_TCCTCAGCCAACCCCCTTTTG |
| *AOS* (AT5G42650) | for_TGGTGGCGAGGTTGTTTGTGATTG |
| | rev_ATTAACGGAGCTTCCTAACGGCGA |
| *AOC2* (AT3G25770) | for_AATTAGATCGACACAGCCCCAAG |
| | rev_CCGAGACCGAACATTAAGCTGA |
| *OPR2* (AT1G76690) | for_AATCGCGGTTTTCAGCCAAG |
| | rev_GCCATTAGCACGCATTTGAG |
| *OPR3* (AT2G06050) | for_TGGTTGGCATGCTCAATAAG |
| | rev_GCCTTCCAGACTCTGTTTGC |
| *DREB2A* (AT5G05410) | for_AGCAACAACAGCAGGATTCG |
| | rev_AGGTCACGTAGAAGCTCATCG |
| *FAD-OXR* (AT4G20860) | for_TTTGGCGTTGTCTTGTCGTG |
| | rev_TGTTCATGCTAGGTCCCATCG |
| *ZAT10* (AT1G27730) | for_AGGCTCTTACATCACCAAGATTAG |
| | rev_TACACTTGTAGCTCAACTTCTCCA |
| *ERF5* (AT2G20880) | for_TGTCGCCGTTATCTCCTCATC |
| | rev_CACGTCAGCATACACATCGTTC |
| *GST6* (At2g47730) | for_AGTCAAGGCAACCACTAACG |
| | rev_TTAAAGACACGCTCGAAGGC |
| *TCH4* (At5g57560) | for_AACGCTGATGATTGGGCAAC |
| | rev_TGAAAGCCACGGTAAGAAGC |
| *JAZ1* (AT1G19180) | for_ATCAACTTGGCGAGCAAAGG |
| | rev_TGCGATAGTAGCGATGTTGC |
| *JAZ2* (AT1G74950) | for_TTTTTCTGCCGAGTGTTGGG |
| | rev_ACCCTTCTCCTTCAGGTAACG |
| *JAZ7* (AT2G34600) | for_TACCCATCTTGAGGCTAACGC |
| | rev_ATCCGAACCGTCTGAACTTCTC |
| *JAZ10* (AT5G13220) | for_AACCAACAACGCTCCTAAGC |
| | rev_TATCTCGGAAACTACGACGG |
| *JAZ13* (AT3G22275) | for_ATAGAGATGGCGAGCAAGGATC |
| | rev_TCCTAACGGTGATTCCAGTCTC |

| | |
|---|---|
| *CLH1* (AT1G19670) | for_GAGTGTAAGGCGACGAAAGC |
| | rev_CATACAACCGGCCATAAACC |
| *NATA1* (AT2G39030) | for_GAGTGTAAGGCGACGAAAGC |
| | rev_CATACAACCGGCCATAAACC |
| *SDI1* (AT5G48850) | for_ACCTTGCGATGTGCCTTATC |
| | rev_AATCATCCGCACCCAAAACC |
| *APR3* (AT4G21990) | for_AAGAGGCTTGGATCGTTGTG |
| | rev_AGCAACCTTCACACCACTTC |
| *ORA59 (AT1G06160)* | for_TCCATGAGAAACCGTCCTAGAG |
| | rev_AATAGGAGGAGGAGGAAGAAGG |
| *YFP* | *for_ACTACAACAGCCACAACGTC* |
| | *rev_TGTTGTGGCGGATCTTGAAG* |

**Supplementary Tab. 7: Primer sequences used for Golden Gate cloning of *OPR3*, *OPR2* and *OPR1* and their subcellular targeting.**

| Template and codon alteration | Primer sequence (5´->3´) |
|---|---|
| OPR3ΔSRL | |
| AtOPR3 CDS(1-205 bp), C205G | for_TTGAAGACAAAATGacggcggcacaagggaactc |
| | rev_TTGAAGACAAacaccatggtgccttcggagatgag |
| AtOPR3 CDS(205-1109 bp), C205G-G1109C | for_TTGAAGACAAgtgtctcccggatccgcaggg |
| | rev_TTGAAGACAAtgcgattatatttattcaactctccatcaatcttga |
| AtOPR3 CDS(1109-1164 bp), G1109C | for_TTGAAGACAAcgcaagacgttttacactcaagatccagttg |
| | rev_TTGAAGACAACGAAccaaaaggagccaagaaaggataatccgtg |
| OPR2 | |
| AtOPR2 CDS(1-1122 bp) | for_TTGAAGACAAAATGgaaatggtaaacgcagaagca |
| | rev_TTGAAGACAACGAAccagctgttgattcaagggaagggta |
| OPR1 | |
| AtOPR1 CDS(1-1122 bp) | for_TTGAAGACAAAATGgaaaacggagaagcaaaacagag |
| | rev_TTGAAGACAACGAAccagctgttgattcgaggaaagg |
| YFP | |
| YFP CDS(4-729 bp) | for_TTGAAGACAATTCGgtgagcaagggcgaggag |
| | rev_TTGAAGACAAAAGCttatctaatagccgcgttnttgtacagc |
| YFP-HDEL | |
| YFP CDS(4-726 bp)-HDEL | for_TTGAAGACAATTCGgtgagcaagggcgaggag |
| | rev_TTGAAGACAAAAGCttatagctcgtcatgcttgtacagctcgtccatgc |
| YFP-SRL | |
| YFP CDS(4-726 bp)-SRL | for_TTGAAGACAATTCGGtgagcaagggcgaggagc |
| | rev_TTGAAGACAAAAGCgaggcgggacttgtacagctcgtccatgcc |
| YFP-SKL | |
| YFP CDS(4-726 bp)-SKL | for_TTGAAGACAATTCGgtgagcaagggcgaggag |
| | rev_TTGAAGACAAAAGCttagagcttggatctaatagccgcgttnttgtacagct cg |
| YFP-NES | |
| YFP CDS(4-726 bp)-NES | for_TTGAAGACAATTCGGtgagcaagggcgaggag |
| | rev_TTGAAGACAAAAGCttaatcaagagtaagtctctcaagcggtggtagctg aagtctaatagccgcgttnttgtacagc |
| RECA cTP | |
| AtRECA CDS (1-51 bp) | for_TTGAAGACAAaatggattcacagctagtcttgtctc |
| | rev_TTGAAGACAAacctctgtagacggtaaccggagaatagag |
| Rieske mTP | |
| SlRieske CDS(1-100 bp) | for_TTGAAGACAAAATGcttcgagtagcaggtagaag |
| | rev_TTGAAGACAAACCTccgctaggatctccaggtg |
| SV40 NLS | |
| SV40 CDS(1-45 bp) | for_TTGAAGACAAAATGcttcgagtagcaggtagaag |
| | rev_TTGAAGACAAACCTccgctaggatctccaggtg |
